# Molecular access control for random-access DNA data storage

**DOI:** 10.64898/2026.08.12.744375

**Authors:** Bas W.A. Bögels, Robin T. Vermathen, Angelina Yurchenko, Christopher N. Takahashi, Albert J. Markvoort, Tom F. A. de Greef

**Author notes:** These authors contributed equally: B.W.A. Bögels, R.T. Vermathen.

## Abstract

DNA data storage offers exceptional density and millennial-scale stability, with advances in encoding schemes and reduced synthesis costs making large-scale archiving increasingly viable. However, while efforts have focused on reliable data retrieval, securing DNA-encoded information against unauthorized access remains largely unexplored. Here, we introduce DNA-GUARD (DNA Gated Unlocking and Access Restriction of Data), a molecular access-control system that restricts PCR-based retrieval and complements data encryption. Chemically modified “locker strands” outcompete PCR primers and, through 3′ inverted dT modifications, block polymerase extension and amplification of sequences required for decoding. Complementary “key strands” tethered to magnetic particles sequester and remove locker strands, restoring access. We demonstrate scalability from 555-byte to 1-MB files, orthogonal control of multiple files in mixed libraries, and reliable repeated locking–unlocking cycles. DNA-GUARD is compatible with established DNA storage workflows and provides reversible physical access control that increases the sequencing effort required for unauthorized retrieval.

## Main text

As global data generation accelerates, it is projected to soon surpass the storage capacity offered by conventional data storage technologies^1^. DNA offers a compelling alternative, with exceptional data density, reaching up to 17 EB/g^2^, and stability over millennia, with half-lives exceeding 1,000 years^3^. Since early demonstrations of synthetic DNA data storage^4^, researchers have developed robust encoding algorithms capable of managing large datasets^3,5–11^, alongside techniques for selective retrieval of DNA-encoded information^8,12–16^.

Despite recent progress, DNA data storage is mainly limited to storage and retrieval only and lacks capabilities to perform more complex operations on the data, a feature that is available in conventional systems. Recently, efforts to incorporate computational functions into DNA storage have emerged^17–19^, however access control that directly gates standard DNA-storage retrieval workflows remains comparatively underexplored. This capability is of paramount importance, particularly for data that includes sensitive or confidential information. A range of molecular-security strategies has been explored, including irreversible molecular kill-switches, DNA nanostructures, chemical modifications, specialised retrieval schemes, hybrid molecular-digital encryption, and programmable molecular systems that conceal information or regulate its accessibility^20–26^. These approaches achieve security through different mechanisms, including irreversible erasure, digital or hybrid encryption, specialized molecular readout, and dedicated molecular architectures, rather than providing a reversible access-control layer directly integrated into standard PCR-based random access^7,8^. One recent method^27^ uses patterns of canonical adenine and non-canonical 2-aminoadenine that cannot be faithfully reproduced by DNA polymerases, causing the encoded information to be irreversibly erased during PCR amplification. This molecular non-replicability provides strong protection against PCR-based copying but differs fundamentally from reversible access control, as amplification cannot faithfully reproduce the protected molecular state. An ideal access control scheme for DNA data storage should reversibly restrict physical access to DNA molecules and be compatible with standard PCR-based random access, enabling easy adaptation to existing DNA data storage solutions. Here, we present DNA Gated Unlocking and Access Restriction of Data (DNA-GUARD), a method that controls access at the physical level in standard PCR-based data retrieval by restricting molecular replication rather than transforming the encoded information. This is a fundamental departure from digital encryption approaches, which protect content but not retrieval.

Most DNA data storage techniques encode digital information onto multiple DNA strands, which can be retrieved through polymerase chain reaction (PCR), sequencing, and decoding (Fig. 1a)^28^. DNA-GUARD controls access by selectively interfering with PCR-based retrieval (Fig. 1b). The system introduces “locker strands”—chemically modified oligonucleotides that outcompete primers during PCR annealing (Fig. 1c). A 3′ inverted dT modification renders the locker strands non-extendable, such that their binding to data-encoding templates suppresses primer extension and amplification^29^. Because PCR exponentially amplifies differences in amplification efficiency, this competition generates a strong amplification bias over successive cycles^30^. Consequently, decoder-critical templates become strongly depleted from the sequencing pool, rendering the file undecodable under standard PCR-based retrieval conditions at conventional sequencing coverage. To restore access, complementary key strands immobilised on magnetic particles via biotin–streptavidin interactions capture the locker strands, enabling their magnetic removal and subsequent amplification of the previously suppressed templates. DNA-GUARD therefore functions as a physical lock-and-key mechanism in which locker strands suppress amplification of target sequences, whereas cognate molecular keys remove the lockers, thereby restoring retrieval. Unlike cryptographic access control, security derives from physical control of the DNA sample and secrecy of the locker-region sequences under conventional retrieval workflows, rather than from computational hardness

**Figure 1.**
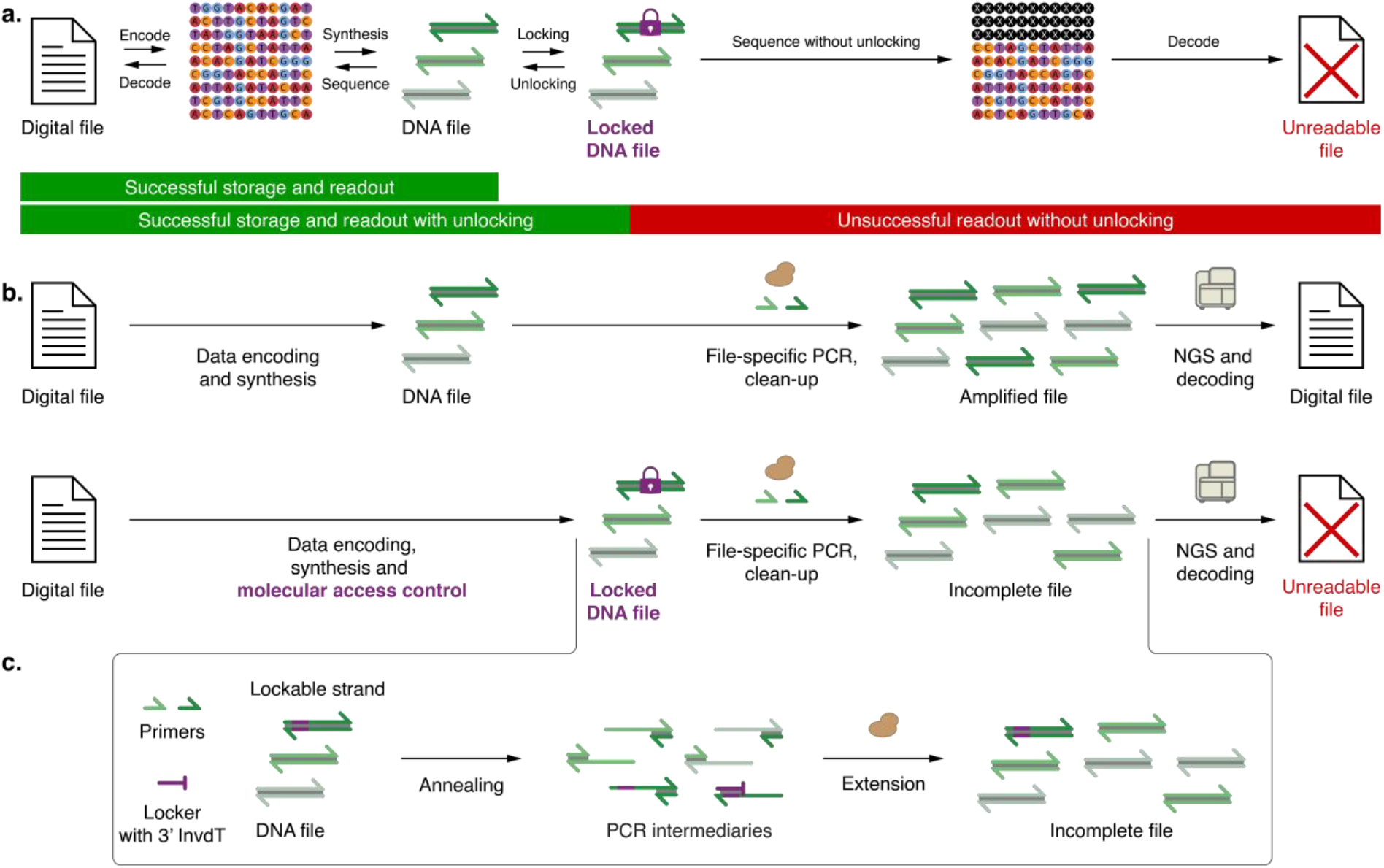
Molecular access control of DNA-encoded digital data. **a**. Cartoon of DNA-based data storage with added access control. A digital file is encoded onto nucleotide sequences represented by their nucleotide sequence. The encoded nucleotide sequence is synthesised as oligonucleotide molecules, yielding the molecularly encoded file. To reverse these steps, the oligonucleotides are sequenced, and the nucleotide sequence is decoded into the original digital file. A locking and unlocking layer is added to the DNA-encoded file by selectively preventing a subset of the oligonucleotides from being accessed. In the locked state, the sequencing of the accessible oligonucleotides yields an incomplete nucleotide sequence, preventing the digital file from being decoded. **b**. Comparison of encoding and retrieval of DNA-encoded digital data with and without access control. Top: Detailed schematic of encoding digital data information into DNA and subsequent retrieval and decoding. A digital file is algorithmically encoded into a set of chemically synthesized DNA strands, yielding a DNA file. To retrieve the encoded file, the strands are amplified using the polymerase chain reaction (PCR) to yield an amplified file. The amplified file is then read using next-generation sequencing (NGS), which allows the digital file to be decoded from the nucleotide sequences. Bottom: Detailed schematic of molecular access control of DNA-encoded data using DNA-GUARD. A digital file is encoded into DNA analogous to that described above. However, a subset of strands that encode the file are in a locked state. Locked strands are strongly depleted during the standard retrieval PCR, resulting in an incomplete representation of the file that is non-decodable at the tested input-read budget. **c**. Reaction diagram of DNA file locking mechanism. The lockable strands within the DNA file contain a common locker region (shown in purple) complementary to the locker strands. During the annealing phase of a PCR cycle, the locker strand outcompetes the forward primer for hybridization with the lockable strands due to their higher binding affinity. Locker strands are modified with a 3’ inverted dT so they cannot be extended by the DNA polymerase, preventing the amplification of locked strands.

We demonstrate that DNA-GUARD enables scalable molecular access control of DNA-encoded data. We first establish a controllable PCR amplification bias in a two-template system, showing that locker strands strongly suppress target amplification, whereas bead-bound complementary keys substantially reduce the induced bias. Secondly, we apply this mechanism to DNA-encoded files to demonstrate functional access control from 555 bytes to nearly 1 MB. Thirdly, we evaluate repeated access with an improved key design, and lastly, test sequence-selective access to three files in a mixed library.

### Selective PCR amplification bias in a two-template system

We established a proof-of-concept system using two double-stranded DNA (dsDNA) templates, to demonstrate molecular access control via selective DNA amplification. Templates A (***A***_***1***_***A***_***2***_) and B (***B***_***1***_***B***_***2***_) share a common primer set (***U***_***FW***_ and ***U***_***RV***_), mimicking a typical DNA file where many unique dsDNA templates are PCR-amplified by universal primers. We designed a non-extendable locker strand ***L***_***1***_ to selectively suppress template A. During PCR annealing, ***L***_***1***_ competes with forward primer ***U***_***FW***_ for binding to strand ***A***_***2***_. A 3’ inverted dT modification blocks DNA polymerase extension, preventing amplification of template A (Fig. 2a). While ***L1*** could partially bind to ***B2***, this interaction did not measurably inhibit amplification (Supplementary Table 2). Template-specific primer sets enabled quantification of each template by quantitative PCR (qPCR; Fig. 2b). We titrated ***L1*** to determine the optimal concentration for selective PCR inhibition. Using an equimolar mixture of templates A and B (40 pM each) and 500 nM primers (***U***_***FW***_ and ***U***_***RV***_), we performed PCR amplification in the presence of 0, 50, 500, or 5000 nM locker strand ***L***_***1***_, then purified the products and measured the relative PCR amplification bias using qPCR (Methods). The difference in Ct values between template A and template B (ΔCt) was compared across locker strand concentrations (Fig. 2c). At 50 nM ***L***_***1***_, we observed a 1.25-cycle increase in ΔCt compared with the 0 nM locker state. At 500 nM ***L***_***1***_, ΔCt increased by 8.26 cycles relative to the no-locker control. Increasing ***L***_***1***_ to 5000 nM produced only a further 0.83-cycle change that was not statistically significant. For subsequent experiments, 400 nM locker strand was selected as a near-maximal concentration within the tested range.

**Figure 2.**
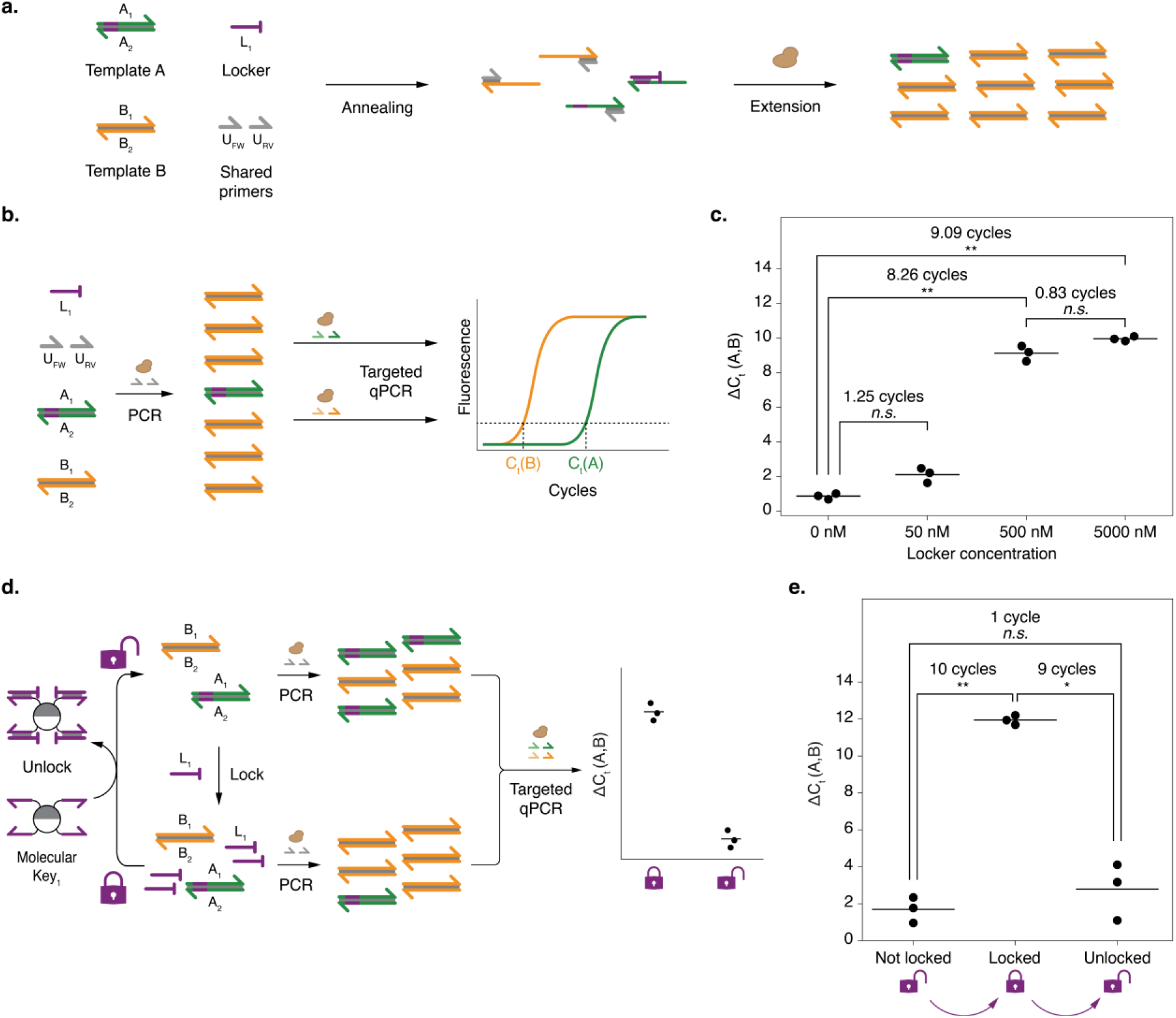
Demonstrating reversible PCR bias in a two-template system. **a**. Reaction diagram of biased amplification in a two-sequence system. Two templates, A (**A**_**1**_**A**_**2**_, green) and B (**B**_**1**_**B**_**2**_, orange) share a set of primers (**U**_**FW**_ and **U**_**RV**_) for PCR-based amplification. Additionally, a region complementary to locker strand **L**_**1**_ is included in the A template (indicated in purple). The shared primers can hybridize with templates A and B during PCR-based amplification. Locker strand **L**_**1**_ hybridizes with **A**_**2**_, thereby preventing the binding of primer **U**_**FW**_. A 3’ inverted dT modification renders **L**_**1**_ non-extendable, preventing amplification of template A. **b**. Cartoon describing induced amplification bias measurement using qPCR. The relative concentration, as expressed in the threshold cycle (C_t_) value, of both templates after biased amplification can be determined using quantitative PCR (qPCR) by employing a primer pair specific for either template A (**F**_**A**_ and **R**_**A**_) or template B (**F**_**B**_ and **R**_**B**_). **c**. Effect of varying locker strand concentration on PCR amplification bias. Solid lines denote the mean ΔCt values (ΔCt = Ct(A) − Ct(B)) from three experiments per locker concentration; points represent individual experiments. Results for all samples containing **L**_**1**_ were compared to the 0 nM **L**_**1**_ condition. A repeated-measures ANOVA followed by post-hoc paired t-tests for all pairwise comparisons with Holm–Bonferroni correction revealed, relative to the 0 nM **L**_**1**_ condition, a 1.25-cycle difference at 50 nM **L**_**1**_ that was not statistically significant (p = 0.08), a statistically significant differences of 8.26 cycles at 500 nM **L**_**1**_ (p = 0.004) and 9.09 cycles at 5000 nM **L**_**1**_ (p = 0.002) and a 0.83-cycle difference between 500 nM and 5000 nM **L**_**1**_ that was not statistically significant (p = 0.08). **d**. Cartoon describing locking and unlocking of a two-template system using molecular keys. As in **b** and **c**, template A can be locked by adding **L**_**1**_. Template A is unlocked by incubating with magnetic bead-bound **Key**_**1**_ and subsequent magnetic removal of magnetic bead-bound **Key**_**1**_**-L**_**1**_ complexes. Unlocked and locked solutions can both be amplified using shared primers **U**_**FW**_ and **U**_**RV**_. The relative amounts of dsDNA template A and B after PCR can be determined as described in **b**.. **e**. Differences in C_t_ values between templates A and B after a locking-unlocking cycle. Solid lines denote the mean ΔCt values for three experiments; the points represent individual experiments. Relative template concentration differences were determined using qPCR as described in **b**.. A repeated-measures ANOVA followed by post-hoc paired t-tests with Holm–Bonferroni correction revealed significant differences in ΔCt between the locked state and the no locker and unlocked states of 10 cycles (p = 0.0018) and 9 cycles (p = 0.023), respectively. Measured C_t_ values are listed in Supplementary Table 3. The sequences for **A**_**1**_, **A**_**2**_, **B**_**1**_, **B**_**2**_, **U**_**FW**_, **U**_**RV**_, **L**_**1**_, **Key**_**1**_, **F**_**A**_, **R**_**A**_, **F**_**B**_ and **R**_**B**_ are provided in Supplementary Table 1. Stars indicate statistical significance: n.s.: not significant (p ≥ 0.05); *: p < 0.05; **: p < 0.01; ***: p < 0.001; ****: p < 0.0001.

Having demonstrated selective PCR bias, we tested whether removing ***L***_***1***_ could reverse the effect. Bi-otinylated key strand ***Key***_***1***_, the reverse complement of ***L***_***1***_, was tethered to streptavidin-coated super-paramagnetic particles, yielding “molecular keys”. These molecular keys sequester ***L***_***1***_ through hybridi-zation, allowing magnetic removal of the complex (Fig. 2d). We tested this unlocking mechanism using three conditions. First, we PCR-amplified the same equimolar mixture of templates A and B used pre-viously (not locked control). Second, we added 400 nM ***L***_***1***_ to the template mixture and directly PCR-amplified it (locked condition). Third, we incubated the locked condition template mixture with mo-lecular keys, subsequently magnetically removed the bead-bound ***Key***_***1***_***-L***_***1***_ complexes (Methods), and then PCR-amplified the solution (unlocked condition). qPCR measurements revealed that locking increased ΔCt from 1.69 ± 0.69 to 11.94 ± 0.25 cycles (mean ± standard deviation). Cognate-key treatment reduced ΔCt to 2.79 ± 1.54 cycles (mean ± standard deviation), a value not significantly different from the no-locker control, demonstrating reversible control of the induced amplification bias.

### Locking DNA-encoded data using molecular access control

Having demonstrated controllable PCR amplification bias into the two-template system, we next applied this approach to DNA-encoded data. We developed an encoder/decoder based on dual Reed-Solomon coding schemes^3,8^ that designates specific sequences for selective amplification control using ***L***_***1***_ (Methods and Supplementary Note 1). As a proof of principle, we encoded a text file (***File 1***; Supplementary Table 4) into 42 DNA sequences, six of which contained a locker (L) region complementary to ***L***_***1***_ (Fig. 3a). In the presence of ***L***_***1***_, PCR preferentially amplifies the 36 non-lockable strands while strongly suppressing amplification of the six lockable strands. The loss of all six lockable strands exceeds the number that the outer code can recover, leaving the file below the decoding threshold (Supplementary Table 5). This selective depletion, therefore, prevents the decoder from reconstructing the complete file. Treatment with the cognate molecular key ***Key***_***1***_, followed by magnetic removal of the key-locker complexes, restores the lockable strands to a level sufficient for decoding. We termed this encoding and access-control workflow DNA-GUARD (DNA-Gated Unlocking and Access Restriction of Data).

**Figure 3.**
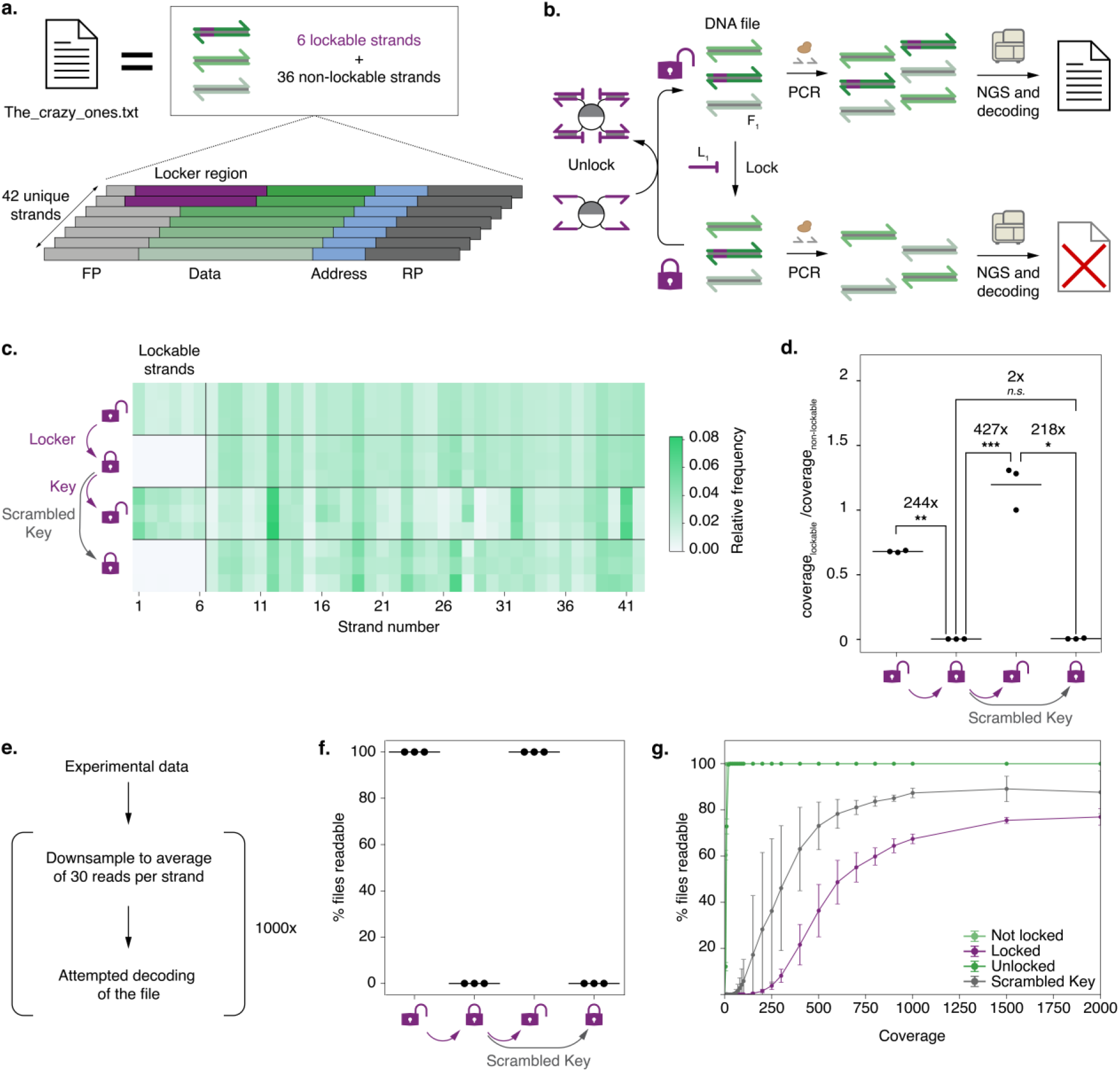
Locking and unlocking of digital data using DNA-GUARD. **a**. Cartoon of data encoding using DNA-GUARD. A 555-byte text file was encoded onto 42 DNA strands (**File 1**). All strands share two universal primer-binding sites, FP and RP, and an internal addressing structure to reorder the data after sequencing. Furthermore, 6 out of the 42 strands contain the additional L region (indicated in purple), allowing these strands to be locked by preventing their amplification. A description of the encoding algorithm is provided in Supplementary Note 1. **b**. Cartoon describing locking and unlocking of DNA-encoded data. **File 1** can be locked to prevent decoding by adding locker strand **L**_**1**_. PCR of the file using universal primers **U**_**FW**_ and **U**_**RV**_ in the presence **L**_**1**_ greatly reduces amplification of lockable strands, which results in an incomplete, and therefore unreadable, file after decoding. To unlock **File 1**, molecular keys, consisting of magnetic bead-bound **Key**_**1**_ strands complementary to **L**_**1**_, are added to the solution. Upon sequestration of **L**_**1**_ by the molecular keys, **L**_**1**_-**mKey**_**1**_ hybrids can be removed using magnetic retrieval. Subsequently, PCR can amplify the unlocked file, and the resulting DNA can be sequenced for file decoding. **c**. Heatmap of relative number of sequencing reads of file-encoding strands after PCR in not locked, locked state, unlocked state with **Key**_**1**_ and after adding scrambled **Key**_**S**_. Both locked and unlocked samples were amplified using PCR before sequencing, and each strand’s relative number of reads was determined (Methods). Each square denotes relative reads for a specific strand in a specific replicate (n = 3). **d**. Ratio of relative **File 1** reads for lockable strands to non-lockable strands in locked and unlocked states. Not locked, locked, unlocked and scrambled Key state **File 1** PCR products were sequenced, and the relative number of reads was determined for each strand (Methods). The ratios of the mean reads of strands that contain the L region to the mean reads of strands that do not contain the L region. A log-scaled repeated measures ANOVA test with post-hoc paired t-tests per pair with Holm-Bonferroni correction revealed statistically significant (p=0.0011) 244-fold difference between the not locked **File 1** solution and locked **File 1**, and a statistically significant (p=0.0001) 427-fold difference between locked and unlocked **File 1**. A statistically not-significant 2-fold difference (p=0.123) was found between the locked state and the sample “unlocked” with scrambled **Key**_**S**_. A statistically significant 218-fold difference (p=0.011) was found between the **Key**_**1**_ unlocked state and the **Key**_**S**_ unlocked state. A statistically significant 125-fold difference (p=0.013) was found between the not locked state and the **Key**_**S**_ unlocked state. A statistically significant (p=0.040) 1.7-fold difference between not locked and unlocked state was measured, this effect is discussed in more detail in figures **4 c** and **d**. Measured mean reads and coverage ratios are listed in Supplementary Table 6. **e**. Simulation used to determine functionality of DNA-GUARD as a data protection method. Reads obtained from PCR of **File 1** are randomly sampled using Monte Carlo downsampling to simulate an average coverage of 30 reads per strand. Decoding is attempted, and the result is recorded. These steps are repeated 1000 times to determine the percentage of times **File 1** can be accessed in each state. **f**. Percentage of tries **File 1** is readable after simulated samplings. The percentage of trials **File 1** could be decoded for each sequenced experiment was determined before locking, in a locked state, after unlocking with **Key**_**1**_ and after addition of **Key**_**S**_. In the unlocked state, **File 1** could be decoded in 100% of the trials; in the locked state or after attempted unlocking by **Key**_**S**_ **File 1** could be decoded in 0% of the trials. Solid lines denote mean percentage of readable trials; points indicate individual experiments. **g**. Evaluating robustness of DNA-GUARD against high sequencing coverage as a means to bypass access control. Downsampling and subsequent decoding attempts as in f were performed for sequencing coverages ranging from 0x to 2000x. Points denote mean percentage of readable trials; whiskers denote standard deviations. Sequences for **U**_**FW**_, **U**_**RV**_ **L**_**1**_, **Key**_**1**_, **and Key**_**S**_ are given in Supplementary Table 1. The original text file encoded onto DNA is shown in Supplementary Table 4. Sequences encoding for **File 1** are made available as stated in the data availability section. Stars indicate statistical significance: n.s.: not significant (p ≥ 0.05); *: p < 0.05; **: p < 0.01; ***: p < 0.001; ****: p < 0.0001.

We quantified DNA-GUARD’s impact on PCR amplification bias using NGS (Fig. 3b). Sequencing confirmed strong depletion of lockable strands in the locked state and restoration following treatment with the cognate molecular key (Fig. 3c, Supplementary Fig. 1). Without locking, no systematic difference in representation is expected between lockable and non-lockable strands. Locked ***File 1*** showed strong depletion of the six lockable strands, demonstrating that the locking mechanism remains effective in a DNA-encoded file. Unlocking restored sequencing coverage of the lockable strands to a level sufficient for decoding. We quantified the relative representation of lockable and non-lockable strands as the ratio of their mean read counts (Fig. 3d). Before locking ***File 1***, this ratio was 0.680 ± 0.008 (mean ± standard deviation), indicating modest baseline underrepresentation of the lockable strands. Locking reduced the ratio approximately 244-fold to 0.0028 ± 0.0004, demonstrating strong selective depletion of the lockable population. Cognate unlocking with ***Key***_***1***_ increased the ratio to 1.20 ± 0.17, restoring the lockable strands to a level sufficient for decoding, whereas treatment with the scrambled key ***Key***_***S***_ left the ratio at 0.006 ± 0.0036, indicating continued suppression. Thus, unlocking restored functional access without requiring the original strand distribution to be reproduced. The failure of the scrambled key to restore the lockable population further demonstrates that DNA-GUARD unlocking is sequence-specific.

Having demonstrated controllable amplification bias, we tested whether DNA-GUARD prevents data access under realistic conditions. At very high initial sequencing coverage, small numbers of lockable reads were detected even when the file was locked, which could theoretically allow file reconstruction. However, typical DNA data storage uses much lower coverage^6,8^. To estimate decoding performance under a controlled read budget, we repeatedly downsampled each independently prepared sample to *N* x *C*quality-filtered reads, where *N* is the number of strands in the file and *C*is the desired average read depth across the file, hereafter referred to as coverage. Each downsampled read set was then passed to the decoding algorithm to assess file decodability. This procedure was repeated 1000 times for each sample to estimate the probability of successful decoding for that specific experimental condition (Methods; Fig. 3e). Throughout this work we typically set *C*= 30, corresponding to an average coverage of 30x, samples unlocked with the cognate key decoded successfully in all trials, whereas locked samples and samples treated with the scrambled key failed to decode in every trial (Fig. 3f). To further assess the robustness of DNA-GUARD, we evaluated all file states across simulated coverages ranging from 0 to 2000× (Fig. 3g). Not locked and unlocked file samples were successfully decoded in 100% of trials at coverages above 10× and 20×, respectively, whereas locked samples and samples treated with ***Key***_***S***_ exceeded 50% decoding success only at coverages of 700× (55 ± 4.5%, mean ± S.D.) and 400× (60 ± 18%, mean ± S.D.), respectively. These results demonstrate that DNA-GUARD substantially increases the sequencing coverage required for file access.

Because DNA-GUARD acts on PCR-based retrieval, we also examined whether direct sequencing of ***File 1*** without prior retrieval PCR could bypass the access-control layer. Direct sequencing did not selectively deplete lockable strands in the locked state (Supplementary Fig. 2a,b). At a simulated coverage of 30×, neither the not locked nor the locked sample could be decoded by direct sequencing (Supplementary Fig. 2c). We next simulated increasing sequencing coverages to determine the depth at which the samples became decodable. The first simulated coverage at which more than 50% of decoding attempts were successful was 250× for the not locked state (60.4 ± 4.7% decodable) and 2000× for the locked state (58.5 ± 27.4% decodable) (Supplementary Fig. 2d). These preliminary results therefore indicate that substantially greater sequencing depth is required for successful direct retrieval of the locked file, although the underlying basis of this difference remains to be established.

### DNA-GUARD scales to megabyte-scale files and supports repeated locking-unlocking cycles

Following the initial demonstration of DNA-GUARD-mediated access control, we tested its robustness along two additional dimensions. First, we examined whether performance was maintained as file size increased from hundreds of bytes to 1 MB. Second, we tested whether DNA-GUARD supported repeated locking-unlocking cycles.

To demonstrate scalability at larger file sizes, we encoded a nearly 1 MB file in DNA (Fig. 4a). The encoded file consisted of 65,419 oligonucleotides—a 1,560-fold increase over the 42 strands used previously—of which 1,200 strands contained the L region marking them as lockable by ***L***_***1***_ (Fig. 4a). We verified that DNA-GUARD is compatible with this increased file size by sequencing locked and unlocked samples and attempting decoding at 30× coverage for 100 trials (Fig. 4b, Supplementary Fig. 3). DNA-GUARD performed effectively at this larger scale, with locked files remaining unreadable and unlocked files decoding successfully in all trials.

**Figure 4.**
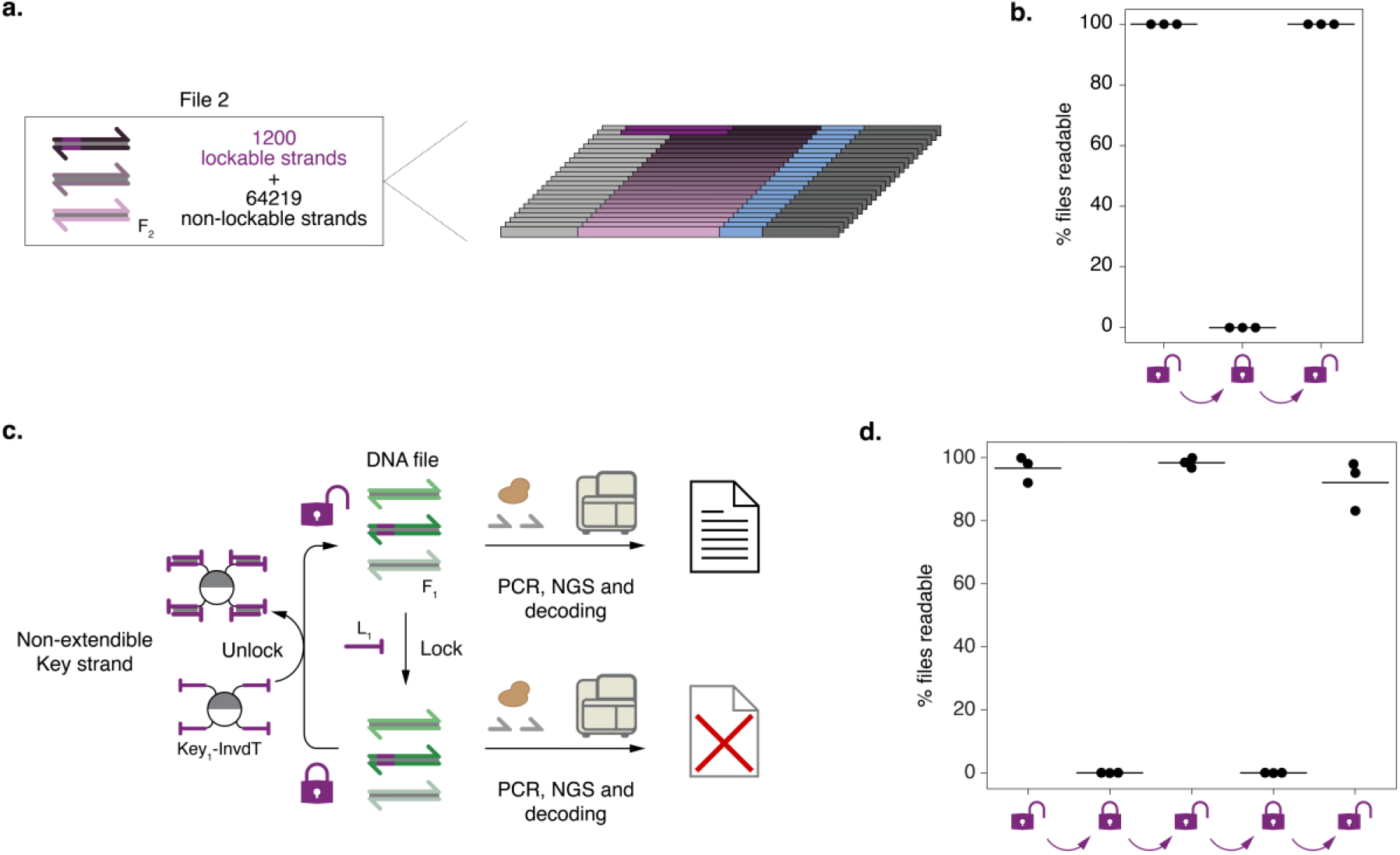
DNA-GUARD scales to megabyte files and supports repeated locking-unlocking cycles. **a**. Cartoon of data encoding for a larger file using DNA-GUARD, analogous to Fig. 3a. A nearly 1-MB file was encoded onto 65,419 DNA strands (**File 2**), of which 1,200 strands contain the L region complementary to **L**_**1**_, marking them as lockable, while the remaining 64,219 strands are non-lockable. **b**. Percentage of tries **File 2** is decodable after 100 simulated samplings at a fixed coverage of 30x. The percentage of trials **File 2** could be decoded for each sequenced experiment was determined before locking, in a locked state, and after unlocking. **c**. Schematic of repeated locking and unlocking of **File 1** using improved DNA-GUARD. The scheme is similar to that of Fig. 3b; however, the molecular key was changed. In the original design, **Key**_**1**_ was only modified to include a 5’ biotin handle for tethering to the magnetic bead. Improved molecular keys using **Key**_**1**_**-InvdT**, a variant of **Key**_**1**_ with an added 3’ inverted dT that prevents it from serving as an unintended primer. **d**. Percentage of tries **File 1** is readable after 1000 simulated samplings at a fixed coverage of 30x before locking, in a locked state, and after unlocking. Locking and unlocking were repeated for a total of two rounds; a slight decrease in performance was observed. All measured readability results are listed in Supplementary Table 8. Sequences for **U**_**FW**_, **U**_**RV**_, **L**_**1**_, **Key**_**1**_ and **Key**_**1**_**-InvdT** are given in Supplementary Table 1. Contents of **File 1** and **File 2** are listed in Supplementary Table 4. Sequences encoding for files are made available as stated in the data availability section.

We next tested whether DNA-GUARD supports repeated locking-unlocking cycles. After two rounds of locking and subsequent unlocking, readability of ***File 1*** decreased to 43.6 ± 15.7% (mean ± standard deviation) across the three replicates and lockable strands became strongly over-represented. (Supplementary Fig. 4, Supplementary Table 7). We hypothesised that key strands detached from the magnetic beads and, because of their complementarity to the lockable templates, acted as unintended PCR primers, thereby compromising subsequent locking. To prevent key strands from acting as unintended primers, we added a 3′-inverted dT modification to the key strands, similar to that of the locker strands. We incorporated this modification into ***Key***_***1***_ to create ***Key***_***1***_***-InvdT*** (Fig. 4c) and tested its performance by repeatedly locking and unlocking ***File 1*** over two rounds, then sequencing the resulting PCR products and attempting to decode them using our simulated 30x coverage pipeline. Unlocking performance remained high across both rounds, with respectively 95.9 ± 4.1 and 91.4 ± 7.8 for the first and second round unlocked samples (Fig. 4d, Supplementary Fig. 5), indicating that free ***Key***_***1***_***-InvdT*** no longer acts as an unintended primer. These results are consistent with 3’ blocking preventing amplification distortion caused by detached, extendable key strands and demonstrate that DNA-GUARD supports reliable repeated access control.

### Orthogonal locker design enables parallel access control of multiple files

Having demonstrated scalability with file size, we next addressed multi-file libraries requiring independent access control. Several methods enable the storage and independent retrieval of many DNA-encoded files from mixed libraries^8,12,13,16^. We therefore designed additional locker sequences compatible with previously developed orthogonal file-retrieval primers, using simulated annealing to iteratively optimize candidate locker sequences (Fig. 5a; Supplementary Note 2). The algorithm minimises four concurrent error terms: deviation from the target Gibbs free energy of locker-template binding, deviation from uniform GC distribution, intramolecular accessibility penalizing sequences that are predicted to form stable intramolecular secondary structures and sequence quality and orthogonality assessed by *k*-mer analysis, which quantifies shared sequence motifs within a candidate sequence and *k*-mers it shares with the lockers and primers already present in the pool, including their reverse complements. Over successive iterations, the combined error function converged as candidate sequences increasingly satisfied all criteria (Fig. 5b). Optimising for both thermodynamic stability and sequence orthogonality yielded three locker-and-primer combinations, these locker-primer combinations showed the expected cognate suppression and recovery in qPCR assays and were therefore advanced to file-level testing (Supplementary Fig. 6).

**Figure 5.**
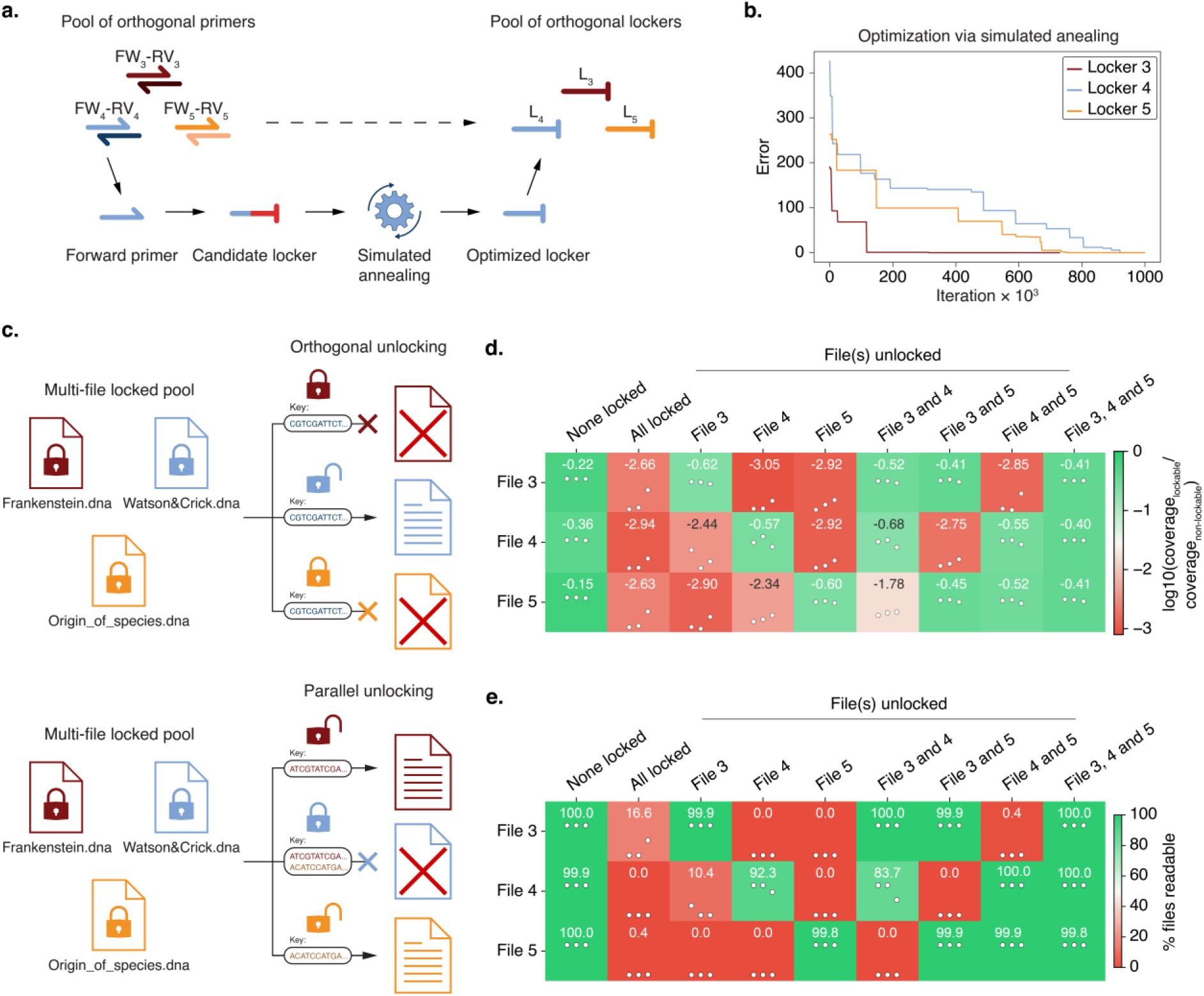
Orthogonal locker design enables parallel access control of multiple files. **a**. Design of orthogonal primer and locker pairs. Starting from prevalidated orthogonal primer pairs, lockers were designed using the 3’ 15 nucleotides of each primer to initialise the locker sequence. Locker sequences were then optimised through a simulated annealing algorithm, yielding a pool of optimised locker sequences. **b**. Multi-objective optimization convergence for de novo locker sequence design. Optimization trajectories for three independent simulated annealing runs to generate **L**_**3**_ (red), **L**_**4**_ (blue) and **L**_**5**_ (or-ange), showing the total error of the sequence with the lowest total error achieved at each iteration. **c**. Schematic of selective unlocking of DNA-encoded files from a mixed pool of files. Three DNA-encoded files and their respective lockers are mixed to form a pool of locked files. The intended file can be unlocked by adding the correct file-specific molecular key, while the other files are unaffected. PCR, sequencing and subsequent decoding of the pool result in only the intended file being readable. **d**. Ratio of relative reads for lockable strands to non-lockable strands for **File 3, File 4** and **File 5** before locking, after all files were locked, and after either single files or multiple files were unlocked. **e**. Measured orthogonality of locking and unlocking in a mixed pool of three files. Percentage of tries **File 3, File 4** or **File 5** is readable after 1000 simulated samplings at a fixed coverage of 30x. The percentage of trials for which files could be decoded for each sequenced experiment was determined before locking, after all files were locked, and after either single files or multiple files were unlocked. Measured values per replicate are listed in Supplementary Table 10 – 12. Sequences for **FW**_**3**_, **RV**_**3**_, **L**_**3**_, and **Key**_**3**_, for **FW**_**4**_, **RV**_**4**_, **L**_**4**_, and **Key**_**4**_ and for **FW**_**5**_, **RV**_**5**_, **L**_**5**_, and **Key**_**5**_ are given in Supplementary Table 9. Content of files encoded onto DNA are shown in Supplementary Table 4. Sequences of **File 3, File 4** and **File 5** made available as stated in the data availability section.

Having established a method for generating orthogonal locker sequences, we tested selective locking and unlocking of multiple files within a mixed library. Because the original primer–locker pair was not designed for orthogonality, we encoded and synthesised three new files ***File 3, File 4*** and ***File 5***, which can be retrieved using the orthogonal primer pairs ***FW***_***3***_***-RV***_***3***_, ***FW***_***4***_***-RV***_***4***_ and ***FW***_***5***_***-RV***_***5***_ and locked using their corresponding lockers ***L***_***3***_, ***L***_***4***_ and ***L***_***5***_ (Supplementary Table 5). Following the same workflow used for ***File 1*** in Fig. 3, all generated locked-key pairs were confirmed to effectively lock and unlock their respective files (Supplementary Fig. 7).

To assess orthogonality, we combined equimolar amounts of the three files into a single library and added all three corresponding locker strands to lock the library. Selective file access was achieved by adding the cognate molecular key (e.g., magnetic beads carrying ***Key***_***4***_), leaving non-target files locked. We tested three unlocking scenarios: *i)* single-file unlocking, *ii)* two-file unlocking, and *iii)* all files unlocked together, using their respective molecular keys. After unlocking, we individually amplified each file with its specific primers, sequenced the products, and attempted to decode them. Across the three files, locking reduced the mean lockable/non-lockable read ratio by a median factor of 403 (interquartile range 86–1045), whereas cognate-key treatment restored a median of 57% of the corresponding baseline (interquartile range 41–69%). At a nominal filtered-input read depth of 30×, all nine baseline file replicates were at least 99.8% readable. Of the 36 file-state replicates intended to remain locked, 34 reached at most 1.3% readability; two were partially readable (49.9% and 31.3%), and both showed unusually weak suppression, the larger occurring in the fully locked state with no key present. The second occurred with a non-cognate key present, but such keys did not measurably change the lockable/non-lockable ratio of locked files (1.01-fold relative to the fully locked state; p = 0.99, paired, n = 9 file–replicate pairs), and in both cases the same locker condition gave 0% readability in the other two replicates, indicating incomplete suppression rather than systematic cross-unlocking. Of the 36 intended-unlocked replicates, 34 were at least 99% readable, whereas two ***File 4*** replicate-3 conditions reached 77.0% and 51.0%; in these, the cognate key released only 21% and 18% of the lockable strands, well below the median of 57%, indicating incomplete unlocking rather than interference by other keys. Together, these observations establish functional orthogonality of the three locker-key systems under the tested workflow.

## Conclusions

Here, we introduce DNA-GUARD, a molecular access-control system for DNA data storage that selectively blocks PCR amplification of decoder-critical strands. Unlike computational encryption, which protects information by transforming its representation, DNA-GUARD restricts molecular replication and thereby prevents standard PCR-based retrieval at conventional coverage in the absence of the cognate molecular key. The system integrates directly with established random-access workflows and enables reversible, orthogonal file locking within mixed DNA libraries. Across the different file sizes and access-control configurations, locker strands strongly depleted decoder-critical strands, whereas cognate keys restored file decodability, with robust discrimination between locked and unlocked states from 555 bytes to nearly 1 MB. In a mixed three-file library, cognate keys enabled selective file access, with only occasional replicate-specific failures. Together, these results establish DNA-GUARD as a reversible molecular access-control strategy that complements rather than replaces digital cryptography.

DNA-GUARD enables file-specific access control within large DNA archives. Importantly, the relative overhead decreases with file size. In the 42-strand proof-of-concept, 14% of strands were lockable, whereas in the nearly 1 MB file containing approximately 65,000 strands, only 1.8% were lockable. This reduced fraction limits the additional synthesis burden to less than 0.5% of the encoded bases. The operational cost per unlocking event is dominated by magnetic bead reagents and remains constant regardless of library size, representing a negligible fraction of the initial synthesis investment (Supplementary Note 3). Importantly, DNA-GUARD offers reversible access control, a key advantage over irreversible molecular “kill-switch” deletion strategies that destroy rather than protect data. Because DNA-GUARD components are DNA oligonucleotides, the security mechanism is expected to match the chemical stability of encoded files, enabling secure archival storage under appropriate conditions. Complementary to our approach, a recent study^31^ demonstrated that DNA can serve as a physical entropy source for one-time-pad key distribution with unconditional security, suggesting that molecular information security is rapidly emerging as a field spanning the full information security landscape from storage to transmission.

DNA-GUARD provides physical access control that complements digital cryptography as part of a defence-in-depth approach. Conventional encryption protects the information content but relies on computational hardness. Once the DNA sequences are recovered, security depends entirely on the strength of the cryptographic scheme and protection of the corresponding keys. DNA-GUARD complements digital cryptography by restricting the standard PCR-based retrieval workflow. As formalised in the security model (Supplementary Note 4), the access-control layer is specific to the PCR-based retrieval explored in this work and may be bypassed by direct sequencing, alternative retrieval strategies, or sufficiently deep sequencing. Preliminary results for File 1 indicate that, in the locked state, direct sequencing without retrieval PCR required substantially greater sequencing depth for successful decoding than the standard PCR-based retrieval workflow (Supplementary Fig. 2). Future work should establish the basis and generality of this effect and explore additional molecular barriers, such as modifications that inhibit adapter ligation or storage at ultra-low copy number, to further impede unauthorised retrieval.

The orthogonal locker-key design naturally extends to multi-party authorisation schemes, in which access to sensitive data requires multiple distinct molecular keys, analogous to cryptographic secret sharing implemented at the molecular level. Practical implementation will require further evaluation of error rates and unlocking kinetics across varied storage conditions. Together with its demonstrated scalability, reversibility, and workflow compatibility, DNA-GUARD provides a foundation for programmable molecular access control in DNA archives, making access itself a property of the molecular retrieval process rather than solely of the encoded information.

## Author Contributions

B.W.A.B. and R.T.V. conceived, designed the study, performed experiments, analysed the data, and wrote the manuscript. A.Y. created code for generating orthogonal locker and primer pairs. A.J.M. analysed the data. C.N.T. wrote the data encoding algorithm and analysed the data. T.F.A.d.G. conceived, designed, and supervised the study; analysed the data; and wrote the manuscript. All authors discussed the results and commented on the manuscript.

## Acknowledgements

This work was supported by a Vici grant from the Dutch Research Council NWO (NWO, 016.176.622) and ERC Consolidator Grant 2020 (AMIGA).

## Materials and methods

### Materials

All chemicals and magnetic particles used were purchased from Sigma, unless otherwise noted. Enzymes were purchased from New England Biolabs, unless otherwise noted. Reagents used for NGS and library preparation were ordered from Illumina. EvaGreen was ordered from Biotium, and KAPA HiFi HotStart PCR Kit was ordered from Roche.

### DNA oligonucleotide synthesis

Separate DNA oligonucleotides and ***File 1*** were purchased from Integrated DNA Technologies (IDT). Modified oligonucleotides were purchased with HPLC purification, and non-modified oligonucleotides were purchased desalted. Stock solutions (100 μM and 10 μM) were prepared in nuclease-free TE buffer (10 mM Tris, 0.1 mM EDTA, pH 7.5; Integrated DNA Technologies) and stored at −30 °C. All other DNA oligonucleotide pools were purchased at Twist Bioscience.

### PCR

The total reaction volume was 25 µL and consisted of KAPA HiFi HotStart, 0.5 µM primers, and 2 µL file solution. Thermocycling was performed in a T100 Touch thermocycler (Bio-Rad). Following initial denaturation (95°C, 3 min), we performed 12 cycles of: denaturation (98°C, 20 s), annealing (65°C, 15 s), and extension (72°C, 15 s), followed by a final extension at 72°C for 30 seconds before cooling down to 4°C. Double-stranded DNA products were purified from the reaction mixture using a Qiagen PCR extraction kit following the manufacturer’s instructions.

### Magnetic removal of locker strands

Molecular keys were prepared by first washing and resuspending 200 µL Streptavidin-coated DynaBeads (ThermoFisher) according to the manufacturer’s instructions for conjugating biotin-labelled DNA. To the washed and resuspended beads 400 µL of 5 µM biotin-labelled DNA was added. The resulting mixture was incubated under gentle agitation for 60 minutes at room temperature. After the DNA was conjugated, excess unbound DNA was removed according to the manufacturer’s instructions. Finally, the beads were resuspended in a final volume of 25 µL to yield the final molecular key solution.

In a typical unlocking experiment, we first prepared a 0.5 nM solution per file (not locked state). To this solution we added the locker strand(s) to yield locked state, final concentration 5 µM per locker. 25 µL of this file mixture was added to molecular keys to yield unlocked state. The resulting mixture was incubated for 60 minutes at room temperature under gentle agitation. After incubation, the mixture was placed on a magnetic separation rack for 3 minutes to separate out the beads, and the supernatant was pipetted to yield an unlocked file solution. At each stage, a 2 µL aliquot was taken for analysis.

For repeated unlocking experiments, a second batch of molecular key solution was produced following the protocol above, with volumes adjusted to 160 µL Streptavidin-coated beads and 360 µL of 5 µM biotin-labelled DNA. The unlocked file solution was then locked again by adding the locker strand to yield a final concentration of 5 µM. From this re-locked solution, 20 µL was added to the molecular keys, after which the unlocking procedure described above was repeated.

### qPCR

Quantitative PCR was performed using CFX96 Touch Real-Time PCR Detection System (Bio-Rad). The total reaction volume was 25 µL and consisted of KAPA HiFi HotStart, 0.5 µM primers, 1x EvaGreen, and 2 µL sample. Initial denaturation was set to 3 minutes at 95°C, and then 40 denaturation cycles at 98°C for 20 seconds, annealing at 65°C for 15 seconds, and extension at 72°C for 15 seconds were performed, followed by a final extension at 72°C for 30 seconds before cooling down to 4°C. Fluorescence was measured during each annealing step. To prevent evaporation, the plate was sealed with a transparent plate sealer. CFX Maestro Software (Bio-Rad) was used to perform baseline correction and calculate threshold cycles (Ct).

### DNA data encoding

Data was encoded into DNA using a dual-layer Reed–Solomon coding strategy that integrates programmable locker prefixes to restrict access. Digital data are split into fixed-size segments and arranged into a strand matrix, where a subset of strands reserve space for locker sequences. Redundancy is introduced by applying an outer code across columns and an inner code within each strand, yielding parity strands that enforce a minimum number of recovered locked strands for successful decoding. Indexed strands are then converted to DNA using a codon mapping that limits homopolymers and maintains GC balance. Decoding clusters reads, corrects individual strands, reorders them by index, reconstructs missing strands with the outer code, and removes locker prefixes to recover the original data. A detailed description of the encoding and decoding algorithm is given in Supplementary Note 1.

### Library preparation and sequencing

Samples were prepared for sequencing following the Illumina TruSeq Nano DNA Library Prep protocol. Briefly, ends were blunted with the End-Repair buffer (ERP2), then purified with Beckman Coulter AMPure XP beads, and an ‘A’ nucleotide was annealed to the 3’ end with A-Tailing Mix (ATL). Ligation was performed using Illumina sequencing adapters from Illumina’s TruSeq DNA CD or IDT for Illumina Unique Dual Indexes kit, with each sample ligated to a unique Illumina index. Finally, the samples were cleaned using Illumina Samples Purification Beads (SPB) and enriched using a variable cycle qPCR until the signal for each sample reached the inflexion point. Final product length and purity were qualified using a QIAgen QIAxcel. Samples were then individually quantified using qPCR and mixed to create an equal-mass library.

A final library was prepared for sequencing by following the Illumina MiSeq Denature and Dilute Libraries Guide. Sequencing libraries were loaded in the Illumina Miseq at 20 pM, including a 30% control spike-in of ligated PhiX genome.

### Alignment of Illumina sequencing data

Basecalling and demultiplexing of sequenced samples was performed using bcl2fastq. The generated FASTQ-files were then aligned against reference sequences using custom Python code. To generate the distribution of reads across files, the sampled reads were subsequently aligned to the reference sequences using Bowtie 2^32^, before the coverage for each sequence was determined using SAMtools^33^. Only primary alignments were retained (secondary and supplementary alignments excluded), and reads with MAPQ ≥ 20 were kept. For each sample, the lockable/non-lockable ratio was calculated as the mean mapped reads per lockable strand divided by the mean mapped reads per non-lockable strand. For Figure 5d, ratios were log10-transformed and cell values show the mean log10 ratio across the three physical replicates (equivalent to the log10 geometric mean); individual replicate values are overlaid.

Read counts per designed strand were obtained from the alignments described above. In each file the first n strands carry the locker region and constitute the lockable set (8, 12 and 9 strands for Files 3, 4 and 5). For every sample, we computed the ratio of the mean read count per lockable strand to the mean read count per non-lockable strand. Because both sets derive from the same sequencing run, sequencing depth cancels in this ratio, which therefore reflects the locking state rather than the read depth.

Two derived quantities are reported. The suppression factor of a locked sample is the not-locked ratio divided by the locked ratio for the same file and replicate. The fraction released by a cognate key is (r − r_locked) / (r_not-locked − r_locked), with r_locked the mean over the locked samples of the same file and replicate; values of 0 and 1 correspond to no release and to release up to the not-locked reference of that file and replicate. Medians and interquartile ranges are reported because both distributions are right-skewed and one sample exceeds the upper bound.

### File-readability analysis

Raw reads were retained only when the median Phred score was at least 25, neither ‘AAAAAA’ nor ‘TTTTTT’ occurred, and the maximum local alignment score to the sequencing adapter or its reverse complement was below 50 using local alignment with the following scores: *match =2, mismatch=-1, gap opening=-4, gap extension=-1*. For a file containing N designed strands, N x D filtered reads were sampled without replacement to simulate nominal input-read depth D. Primer motifs were then identified in the first or last 30 sequenced bases, and the intervening payload was extracted. Payloads were clustered using Starcode^34^. Full-length clusters were passed to the file-specific decoder. Each physical sample was resampled 100 or 1000 times as stated; the reported readability is the percentage of computational samplings that decoded successfully.

### Orthogonal locker design

We designed orthogonal locker-primer sets using a simulated annealing algorithm adapted from van Roekel et al.^35^ to satisfy specific thermodynamic and orthogonality constraints. Primer sequences were taken from the prevalidated orthogonal set reported previously^8^. Each locker strand contains a 15-nucleotide domain matching the 3’ end of its corresponding primer and a 29-nucleotide enrichment region that increases binding affinity. The optimization algorithm minimizes four concurrent error functions: (i) deviation from a target difference in Gibbs free energy between locker–template and primer–template duplexes, calculated using NUPACK^36^ at 65 °C; (ii) a *k*-mer-based sequence-quality and orthogonality term that penalizes homopolymer motifs, internal sequence redundancy, self-complementary motifs, and sequence motifs shared between lockers; (iii) a compositional term that penalizes local GC-content imbalance and extended GC-rich stretches; and (iv) a hairpin-stability term that penalizes stable intramolecular secondary structures predicted at 20 °C.

The simulated annealing procedure iteratively mutates the enrichment region, accepting mutations that decrease the combined error function or, with temperature-dependent probability, mutations that increase error to escape local minima. Detailed design criteria, equations, convergence analysis, and Hamming distance validation are provided in Supplementary Note 2. The software is made available as stated in the Code and Data Availability section.

### Statistics

All results reporting statistical values were from independent triplicates. Analysis was performed using Python’s SciPy^37^ or Statsmodels^38^ (Python 3.12.2, SciPy version 1.14.1, Statsmodels version 0.14.6) libraries. Statistical significance between more than two paired samples was determined using repeated measures Analysis of Variance (ANOVA) followed by post-hoc analysis, using paired t-tests per pair with Holm-Bonferroni correction. Lockable/non-lockable ratios were log10-transformed before repeated-measures ANOVA. Only values of p<0.05 were considered statistically significant.

The effect of a non-cognate key on a locked file was assessed with a single paired t-test on log10-transformed lockable/non-lockable ratios. For each file and replicate, the median ratio across the three locked states in which another file’s key was present was paired with the ratio of the fully locked state of that same file and replicate, giving nine file–replicate pairs. The reported fold change is the back-transformed mean of the paired differences.

## Code Availability

The following code, along with raw sequencing data will be made available to reproduce key parts of our research and can be used to validate the performance of DNA-GUARD:

- Coverage extraction and analysis pipeline
- Decoding of DNA files and subsampling analysis pipeline
- Scripts for the design of orthogonal locker sequences: https://github.com/angyurchenko/DNA_data_storage

Sequencing data, as well as DNA encoded files, and analysis scripts have been deposited on Zenodo (DOI: 10.5281/zenodo.21890757).

