## Supplementary Information for "Molecular access control for random-access DNA data storage"

### Supplementary Notes

#### *Supplementary Note 1 – Locker-compatible encoding and decoding algorithms*

**Definitions**

$N_T$: total number of strands; $N_L$: number of strands that carry a locker prefix; $N_R$: number of redundant strands; $L$: locker prefix length (bytes); B: payload length per strand (bytes); $RS(n,k)_q$: Reed-Solomon code with alphabet size q, block length n, and message length k.

We adapt a dual Reed–Solomon (RS) concatenated code, similar to those previously applied to DNA storage[1,2] with modifications to insert locker sequences at predefined positions in the final strands. Our encoding algorithm begins by splitting the data to be encoded into chunks of B bytes, where B follows from the number of strands: an index of $\lceil \log_2(N_T) \rceil$ bits is stored alongside the payload within the 130 bits available per strand, leaving B = 14 bytes for File 2 and B = 15 bytes for Files 3–5. For strands 1…$N_L$, we reserve $L$ bytes for a locker sequence prefix by reducing the payload from B to (B–$L$). The chunks are padded to a multiple of B bytes and arranged into an $(N_T - N_R) \times B$ byte matrix, where each row corresponds to one prospective strand and each column aggregates the same byte position across strands. An outer RS code is applied column-wise, extending the number of rows from $N_T$ - $N_R$ to $N_T$, where the outer alphabet is $GF(2^8)$ for files with at most 255 strands and $GF(2^{16})$ otherwise. As the tolerated loss of strands is $N_R$, we require $N_R < N_L$ to ensure that the data cannot be decoded without recovering at least $N_L$-$N_R$ locked strands. Next, an index is added to each row to allow reconstruction of the matrix. To each row of the matrix, we apply the "inner" RS(31,27) code over GF(32), with the message restricted to 26 information symbols, giving 130 bits per strand for payload and index. Each encoded row is 31 symbols and is converted into a DNA sequence of 93 nt using a codon table that prevents long homopolymers (>2 repeats) or imbalanced GC-content.

Decoding reverses these steps, with an additional initial clustering stage:

1. Cluster reads by sequence similarity
2. Decode inner RS on each strand to recover index + payload
3. Reorder strands by index; mark missing strands
4. Apply outer RS to reconstruct missing strands and recover the original matrix
5. Remove locker prefixes and concatenate payloads

File 1 was encoded with an earlier version of the codec, where the payload is 16 bytes per strand with a one-byte index, and the outer RS code over $GF(2^{16})$ is applied to the concatenated payload of the entire file as a single codeword rather than column-wise. For the decoder to reconstruct the file, at least two of the six lockable strands must be recovered (Supplementary Table 5).

***Supplementary Note 2 – Automated locker design algorithm***

***General approach***

Our molecular access control method relies on a sequence-design algorithm that generates a library of orthogonal lockers and their keys (reverse complements of locker sequences), each with thermodynamically balanced locking properties.

We extended the algorithm of van Roekel *et al.*[3] to satisfy the specific thermodynamic and orthogonality constraints of the molecular access control system. Primer design itself lies beyond the scope of this work. We therefore employed a prevalidated pool of orthogonal primers reported by Organick *et al.*[2]. Each locker strand has two distinct functional domains. The first domain (15 nucleotides at the 5′ end) shares sequence identity with the 3′-terminal 15 nt of the primer. The second domain, consisting of the remaining 29 3′ nucleotides, is the enrichment region, which increases the binding affinity of the locker strand to the template compared to that of the primer. The higher binding affinity of the locker means that it outcompetes the primer for template binding, thereby preventing PCR amplification[4].

Our locker design algorithm optimizes sequences within a simulated-annealing framework that combines four error terms into a single objective function (Eq. S.6). First, each locker must achieve the target Gibbs free energy to ensure competitive binding over its corresponding primer (Eq. S.1–S.2). Second, a composite k-mer-based error term promotes sequence orthogonality by penalizing shared 3-mer motifs between sequences, which serve as a proxy for potential cross-hybridization, while simultaneously penalizing homopolymer motifs that compromise sequencing fidelity (Eq. S.3). Third, compositional constraints enforce balanced GC content and limit long GC stretches to promote robust experimental performance (Eq. S.4). Fourth, an explicit hairpin stability penalty, evaluated at the key-mediated retrieval temperature (20°C), ensures that individual locker strands remain structurally accessible (Eq. S.5). Each criterion is described in detail below.

***Thermodynamic stability (ΔG error)***

Each locker must bind its template more strongly than the corresponding primer to prevent PCR amplification[4]. The ΔG error quantifies the deviation from a target binding-energy difference between the locker–template and primer–template duplexes. This difference is defined as:

$$\Delta\Delta G_{\text{candidate}} = \Delta G_{\text{locker–template}} - \Delta G_{\text{primer–template}} \ (S.1)$$

where each $\Delta G$ term is the hybridization free energy predicted by NUPACK 4.0.0.28 [5] at 65 °C (1.0 M $Na^+$, 0.0 M $Mg^{2+}$).

For sequence design, candidate lockers are iteratively optimised by minimising the deviation from the target $\Delta\Delta G$. The target value ($\Delta\Delta G_{\text{target}} = -15.19\ kcal/mol$) was determined from the NUPACK-predicted $\Delta\Delta G$ of a wet–lab–validated functional locker sequence. The $\Delta G$ error is defined as:

$$E_{\Delta G} = \left(\Delta\Delta G_{\text{target}} - \Delta\Delta G_{\text{candidate}}\right)^2 \ (S.2)$$

***Sequence quality and orthogonality (k-mer error)***

Each locker sequence must be internally non-repetitive and distinct from all other sequences in the pool to avoid sequencing artifacts and cross-hybridization. The k-mer error is a composite heuristic that addresses these concerns by screening for repeated k-mers within a candidate sequence and k-mers it shares with the locker and primer pool, along with their reverse complements. The error is defined as:

$$E_{\text{kmer}} = [v(RB_i + GC_i) + RK_i + RKs_c]^2 \qquad (S.3)$$

The first two terms promote sequencing fidelity. $RB_i$ counts A/T homopolymer k-mers (k = 3) that appear more than once in the sequence, penalizing only repeated motifs. $GC_i$ counts all occurrences of G/C homopolymer k-mers (k = 3), including single instances, imposing a stricter constraint due to their propensity to form G-quadruplex structures and their association with sequencing errors. The weighting factor v = 2, as reported by van Roekel et al.[3], further amplifies the contribution of these homopolymer terms.

$RK_i$ counts non-homopolymer k-mers (k = 5) that appear more than once within the sequence, penalizing internal sequence redundancy. $RKs_c$ counts non-homopolymer 3-mers from the candidate sequence that also appear in its own reverse complement or in any existing sequence (or reverse complement thereof) in the pool. This term jointly penalizes self-complementary motifs that could promote intramolecular folding and shared motifs between sequences that could lead to cross-hybridization.

The overlap region of each locker is derived directly from its corresponding primer and cannot be altered. The k-mer error is therefore evaluated over the variable portion of the sequence only. Specifically, intra-sequence terms ($RB_i$, $GC_i$, $RK_i$) are computed over the full sequence minus the contribution of the overlap region alone, while inter-sequence and self-reverse-complement terms ($RKs_c$) are computed over a segment comprising the last (k − 1) nucleotides of the overlap region concatenated with the enrichment region, capturing potential cross-interactions at the junction between the fixed and variable domains.

***Compositional balance (GC error)***

Uniform GC content promotes consistent hybridization and reliable sequencing. The GC error penalizes compositional imbalances that could cause amplification bias or poor sequencing performance. The error is defined as:

$$E_{GC} = E_{ratio} + E_{stretch} \quad (S.4)$$

$E_{ratio}$ penalizes deviations in GC content using a 10-nucleotide sliding window. For each window, the GC count must fall between 3 and 7 (i.e., 30–70%). Windows outside this range incur a penalty equal to the distance from the nearest boundary (e.g., a window with 2 GC bases contributes 1, a window with 9 contributes 2). $E_{stretch}$ counts the number of 5-nucleotide windows composed entirely of G and C bases, effectively limiting the maximum tolerated GC stretch to four consecutive bases.

**Intramolecular accessibility (hairpin error)**

Locker strands must remain structurally accessible for key-mediated retrieval. The hairpin error penalizes candidate sequences that are predicted to form stable intramolecular secondary structures. For each candidate locker, the minimum free energy (MFE) of intramolecular folding is predicted using NUPACK 4.0.0.28[6] at 20°C (1.0 M $Na^+$, 0.0 M $Mg^{2+}$). The error is defined as:

$$E_{hairpin} = \begin{cases} (\Delta G_{MFE})^2 & if\ \Delta G_{MFE} < -2.0 kcal/mol \\ 0 & otherwise \end{cases} \qquad (S.5)$$

Candidate sequences with a predicted MFE more negative than -2.0 kcal/mol are penalized in proportion to the square of their MFE, while sequences with weak or no predicted hairpin structure incur no penalty. The check is performed at 20°C, corresponding to the temperature at which locker strands are retrieved by their complementary key sequences, rather than at the PCR annealing temperature (65°C), as this represents the more stringent condition for hairpin formation.

***Implementation details of simulated annealing***

The four error terms (Eqs. S.2-S.5) must be balanced simultaneously during optimization. We combine them into a single objective function minimized via simulated annealing:

$$E = E_{\Delta G} + \lambda_1 E_{kmer} + \lambda_2 E_{GC} + \lambda_3 E_{hairpin} \qquad (S.6)$$

We set $\lambda_1 = \lambda_2 = \lambda_3 = 10$. The value for $\lambda_1$ follows van Roekel et al.[1], and the same weighting was adopted for the two additional terms.
The optimization uses a geometric cooling schedule where temperature decrease from $T_{start} = 5000$ to $T_{end} = 5$ over $N_{total} = 10^6$ iteration according to $T(i) = T_{start} * f^i$, where $f = \left(\frac{T_{end}}{T_{start}}\right)^{\frac{1}{N_{total}}}$
The simulated annealing algorithm is initialized by generating a random enrichment region sequence, which is then concatenated to the 3' end of the overlap region from the corresponding primer sequence (Figure N2.1). The total error function (Eq. S.6) is calculated for this initial sequence, and the value is set as both the minimum error ($E_{min}$) and reference error ($E_{ref}$). The enrichment region then undergoes iterative random point mutations. After each mutation, the error function is recalculated ($E_{new}$). Mutations that decrease the error ($E_{new} < E_{ref}$) are accepted, and $E_{ref}$ is updated to $E_{new}$ for the next iteration. Whenever $E_{new} < E_{min}$, the minimum error is updated ($E_{min} = E_{new}$), and the corresponding sequence is stored. Mutations that increase the error ($E_{new} \geq E_{ref}$) are accepted with probability $P_{acc} = \min(1, \exp(-\Delta E/T))$, where $\Delta E = E_{new} - E_{ref}$ is the error difference, and T is the optimization temperature. If such mutations are accepted, $E_{ref}$ is likewise updated to $E_{new}$. Higher temperature values (T) increase the probability of accepting unfavorable mutations, enabling the algorithm to escape local minima and explore the sequence space more thoroughly. The optimization continues until either the maximum number of iterations ($N_{total}$) is reached or the minimum error threshold ($E_{min} < 10^{-4}$) is achieved, at which point the algorithm terminates.

Throughout optimization, the sequence corresponding to $E_{min}$ is retained as the best-found solution. Because simulated annealing accepts uphill moves, the current sequence at termination is not necessarily the best encountered. The algorithm therefore tracks and returns the sequence with the lowest error across all iterations.

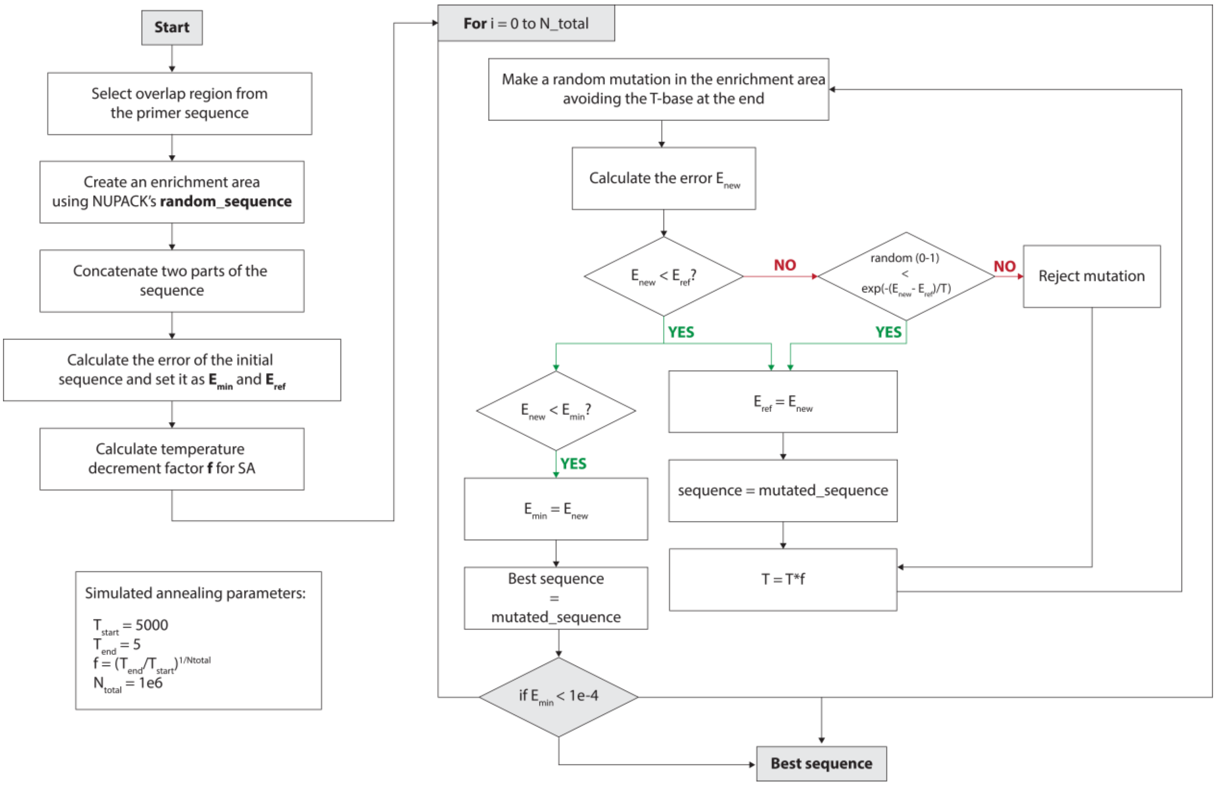

***Figure N2.1 Simulated annealing optimization workflow.*** *Starting from a random sequence, the algorithm iteratively introduces mutations and evaluates them using the combined error function Eq. S.6 comprising four terms: Gibbs free energy deviation ($E_{\Delta G}$), k-mer-based sequence quality and*

*orthogonality ($E_{kmer}$), compositional constraints ($E_{GC}$), and hairpin stability ($E_{hairpin}$). Temperature T decreases over iterations (following a geometrical cooling schedule, with $T_{start}$ = 5000 and $T_{end}$ = 5). Mutations are accepted deterministically if they reduce the error or probabilistically via the Metropolis criterion with temperature-dependent acceptance. The best sequence encountered across all iterations is retained as the final output.*

To generate an orthogonal pool of lockers, each locker sequence was generated sequentially, and the corresponding error functions (Eq. S2-S6) values for the best sequence were recorded for quality analysis. Hamming distances[7] between the resulting sequences were calculated as an additional quality control metric to assess sequence orthogonality. The optimization convergence for the best sequence, including progression of the error terms (Eq S.2-S.6), as well as pairwise Hamming distance comparisons for the generated locker sequences, are presented in Figure N2.2 and Figure N2.3, respectively.

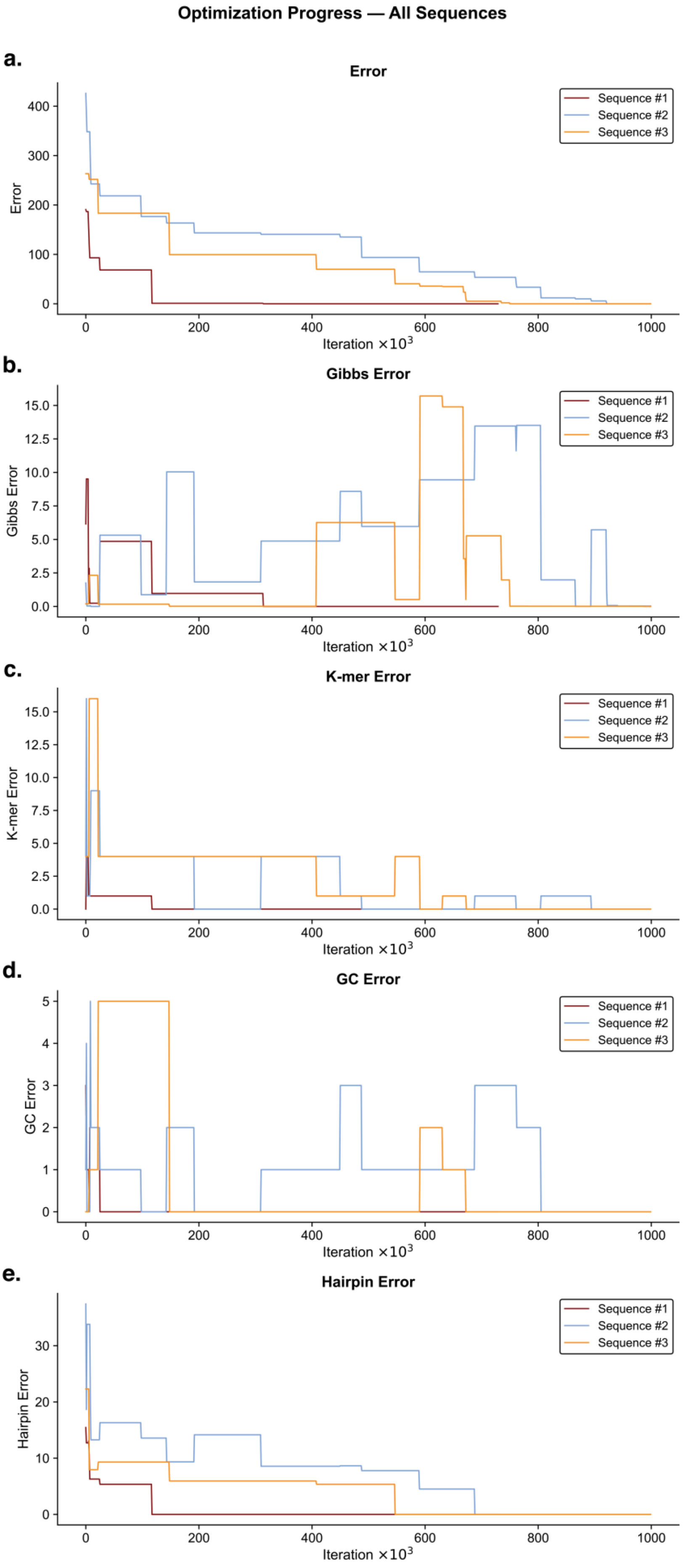

Optimization Progress — All Sequences
a.
Error
Sequence #1
Sequence #2
Sequence #3
Error
Iteration ×10^3
b.
Gibbs Error
Gibbs Error
Iteration ×10^3
c.
K-mer Error
K-mer Error
Iteration ×10^3
d.
GC Error
GC Error
Iteration ×10^3
e.
Hairpin Error
Hairpin Error
Iteration ×10^3

***Figure N2.2 Multi-objective optimization convergence for de novo locker sequence design.*** *Optimization trajectories for three independent simulated annealing runs, showing the error contributions of the best sequence found at each iteration (i.e., the sequence with the lowest total error achieved so far).* ***a.*** *Total Error represents the composite objective function (Eq. S6), combining all weighted penalty terms.* ***b.*** *Gibbs Error ($E_{\Delta G}$) penalizes deviations from the target thermodynamic stability of −15.19 kcal/mol.* ***c.*** *k-mer Error (Ekmer) penalizes repetitive motifs and similarity to existing sequences to enforce sequence diversity and orthogonality.* ***d***. *GC Error (EGC) maintains nucleotide compositional balance within target ranges.* ***e.*** *Hairpin Error (Ehairpin) minimizes undesired secondary structure formation. All three sequences show progressive improvement with Sequence #1 (maroon) and #3 (orange) achieving near-optimal solutions by ~600k iterations, while Sequence #2 (blue) exhibits slower but steady convergence, demonstrating the stochastic nature of the optimization landscape.*

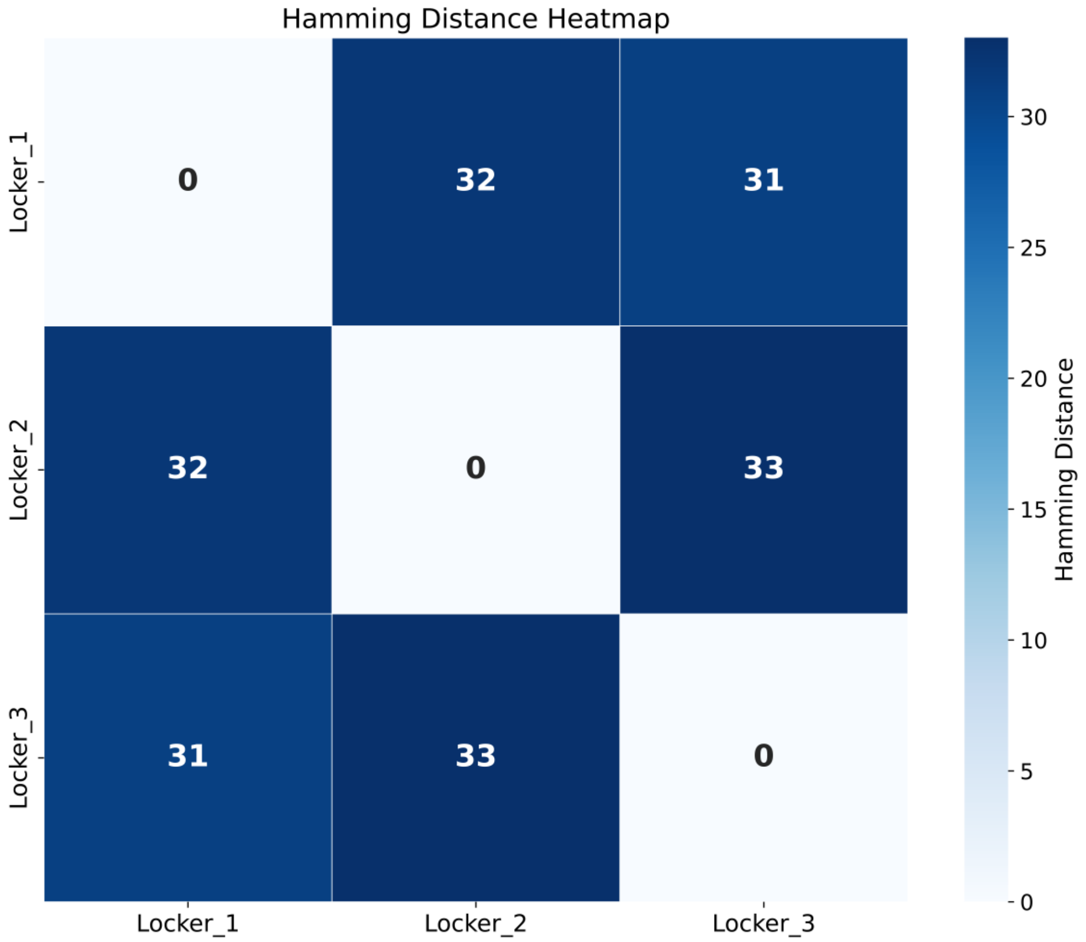

***Figure N2.3 Hamming distance matrix showing pairwise sequence divergence between three optimized locker sequences (45 nucleotides each)****. Values represent the number of differing nucleotide positions between each sequence pair, ranging from 31 to 33 nucleotides (68–73% divergence). This high sequence diversity minimizes the risk of cross-hybridization between different locker sequences during multiplex assays.*

***Supplementary Note 3 – Cost Analysis for DNA-GUARD***

**Overview**

This note provides an order-of-magnitude cost analysis for DNA-GUARD based on vendor list prices (as of October 31, 2025) and the experimental protocols described in the Methods section. We analyse both one-time synthesis costs and per-unlock operational costs, demonstrating that DNA-GUARD adds minimal overhead to DNA data storage systems.

**Assumptions**

*File Characteristics*

The cost structure depends on the number of DNA strands in the stored file and what fraction of them must be lockable to enforce access control: A 1 MB DNA file contains roughly 65,000 strands, of which about 1.8% (≈1,200 strands) incorporate lockable regions. Our small proof-of-concept file (42 strands) used a much higher fraction—14%—because of limited total sequence diversity. The average strand length is 133 nucleotides.

**Experimental Steps Relevant to Costs**

The locking and unlocking steps use several reagents in controlled quantities:

- Locking PCR uses 500 nM of a locker strand in a 25 µL reaction.
- Unlocking uses streptavidin-coated magnetic beads preloaded with a biotinylated key sequence. Each unlocking event starts using 200 µL of bead stock.

The quantities used here directly determine the cost per operation.

**Vendor Pricing for Key Materials**

Costs are estimated using standard prices for modified oligonucleotides and magnetic beads.

*Modified oligonucleotides (Integrated DNA Technologies, October 2025)*

- Base oligonucleotide synthesis: \$0.39/base (25 nmol scale); \$0.67/base (100 nmol scale)
- 5′ Biotin modification: \$46,76 (25 nmol); \$52.47 (100 nmol)
- 3′ Inverted dT modification: \$57.03 (100 nmol)
- HPLC purification: \$46.76 per oligonucleotide (100 nmol scale)

*Magnetic beads*

- Thermo Fisher Dynabeads M-270 Streptavidin:
  - 2 mL \$743.65 list price, price per mL: \$371.825
  - 10 mL \$2,350.65 list price, price per mL: \$235.07

Beads represent the dominant cost in per-unlock operations.

**Synthesis Overhead from Lockable Strands**

Lockable strands include a special region complementary to the key. If this region adds extra nucleotides, the synthesis cost rises proportionally:

$$\text{Overhead} = p_L \times \left(\frac{\Delta L}{L}\right)$$

For the 1 MB file ($p_L$ = 0.018, L = 133 nt):

- ΔL = 0 nt (substitutive design): 0.00% overhead
- ΔL = 20 nt (additive design): 0.27% overhead
- ΔL = 30 nt (additive design): 0.41% overhead

For the 42-strand proof-of-concept (pL = 0.14, L = 133 nt):

- ΔL = 20 nt: 2.1% overhead
- ΔL = 30 nt: 3.2% overhead

Conclusion: At archive scale (pL ≈ 2%), synthesis overhead is negligible (< 0.5%) regardless of L region design. If the L region substitutes for the existing sequence rather than adding length, overhead is effectively zero.

**One-Time Oligonucleotide Purchasing**
Here we calculate the costs of obtaining the modified strands used in locking and unlocking.
*Locker Strand (L1)*
Example: 43 nt strand + inverted dT, HPLC purified (100 nmol scale)

| | | |
|---|---|---|
| Base synthesis: 43 × \$0.67 | \$ | 28.81 |
| 3′ inverted dT modification | \$ | 57.03 |
| HPLC | \$ | 46.76 |
| Total | \$ | 132.60 |

At 400 nM concentration per 25 µL reaction each locking PCR consumes 10 pmol. 100 nmol provides 10,000 locking reactions, so the cost per locking operation: ≈ \$0.013 (1.3 cents), locking is therefore extremely cheap.

*Key Strand (Key1-InvdT)*
Example: 44 nt + 5′ biotin + 3′ inverted dT, HPLC purification (100 nmol scale)

| | | |
|---|---|---|
| Base synthesis: 44 × \$0.67 | \$ | 29.48 |
| 3′ inverted dT modification | \$ | 57.03 |
| 5′ biotin modification | \$ | 52.47 |
| HPLC | \$ | 46.76 |
| Total | \$ | 185.74 |

Price per nmol: \$1.86

**Per-Unlock Operational Costs**
Unlocking requires key DNA and magnetic beads in quantities that dominate recurring expenses. The locker strand cost is negligible in this analysis.

*Key DNA Cost per Unlock*
Each unlocking event uses 2 nmol of biotinylated key:
2 nmol × \$1.86/nmol = \$3.72

*Magnetic Bead Cost per Unlock*
Thermo Fisher Dynabeads M-270 Streptavidin:

- 2 mL \$743.65 list price, price per mL: \$371.825
- 10 mL \$2,350.65 list price, price per mL: \$235.07

Per unlock 200 µL is used, for a minimal cost of \$47.01

**Total Cost per Unlock**

| | | |
|---|---|---|
| Key DNA | \$ | 3.72 |
| Magnetic beads (minimal price) | \$ | 47.01 |
| Locker strand | \$ | 0.01 |
| Total | \$ | 50.74 |

Cost per locking operation: approximately \$0.02; unlocking is clearly the cost-driver.

**Conclusion**
Overall, this cost analysis indicates that DNA-GUARD imposes only a marginal economic burden in relation to the scale of DNA data storage. Locking operations are effectively cost-free, and even unlocking—dominated by bead expenditure—remains inexpensive in absolute terms. It is also

reasonable to expect the prices could be decreased further by buying magnetic beads at scale or moving from streptavidin-biotin-based tethering to cheaper alternatives; however, at present, we cannot provide this analysis. At archive scale, synthesis overhead is negligible, and recurring costs scale strictly with access frequency rather than library size. As a result, DNA-GUARD provides a robust security layer with minimal financial impact, making it well-suited for long-term, infrequently accessed DNA archives.

***Supplementary Note 4 – Security model***

The guarantees of DNA-GUARD should be understood in the context of an explicit adversary model. The protected asset is the digital file encoded across the DNA pool. The legitimate user holds the physical archive, the cognate molecular keys, and knowledge of which strands are lockable and of the locker-region sequences. We consider an adversary who obtains a copy of the DNA archive but possesses neither the cognate keys nor the locker-region sequences, and who attempts retrieval using the conventional random-access pipeline using either universal or file-specific primers, PCR amplification, and next-generation sequencing at typical coverage (≈10–50×).

Against this adversary, locking denies decoding: recovering a locked file requires either the cognate key or sequencing at a coverage roughly one-to-two orders of magnitude above the conventional budget (>700× for PCR-based retrieval and >2,000× for direct sequencing of File 1; Fig. 3g, Supplementary Fig. 2). DNA-GUARD is therefore not a confidentiality mechanism in the cryptographic sense: it raises the sequencing effort required for unauthorized retrieval by a bounded factor and keeps data inaccessible only while the locker-region sequences remain secret and the adversary operates within a conventional read budget. It does not protect against an adversary who knows the locker sequences, who sequences to arbitrary depth, or who bypasses PCR entirely.

For this reason, DNA-GUARD is intended as a physical access-control layer that complements, rather than replaces, encryption of the encoded payload: in a defence-in-depth deployment, payload encryption preserves confidentiality even if the locking layer is bypassed, while DNA-GUARD gates routine retrieval and enables reversible, key-dependent access at the molecular level.

### Supplementary Figures

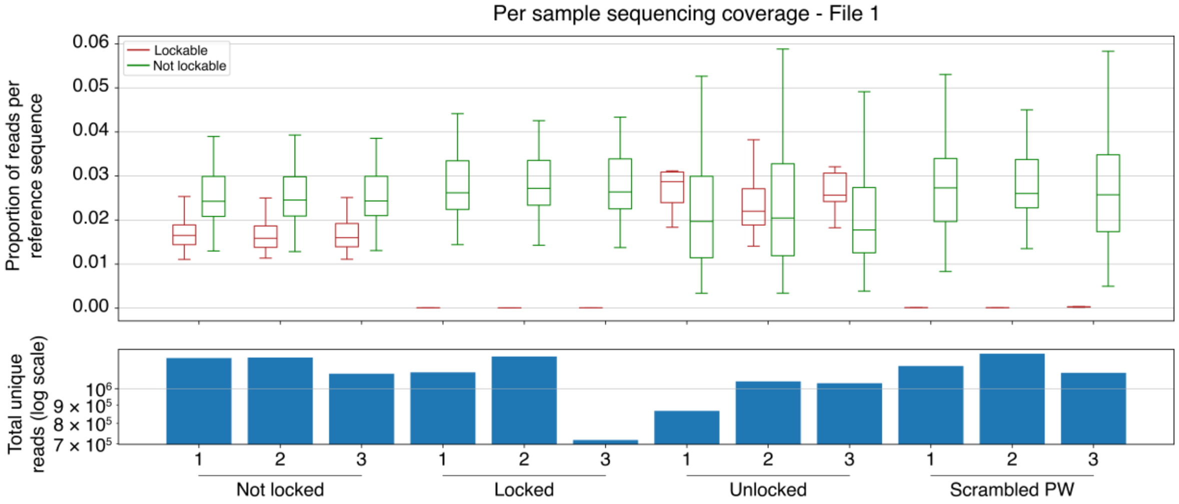

***Supplementary Figure 1 Coverage distributions for each experiment in Figure 3*** *Sequencing coverage across independent experiments for each file state (Not locked, Locked, Unlocked and Scrambled Key). Top: Boxplots indicate coverage distribution for reads that were uniquely mapped to a single reference sequence. Lockable and non-lockable strands are shown as individual populations. Bottom: Total number of aligned reads per sample.*

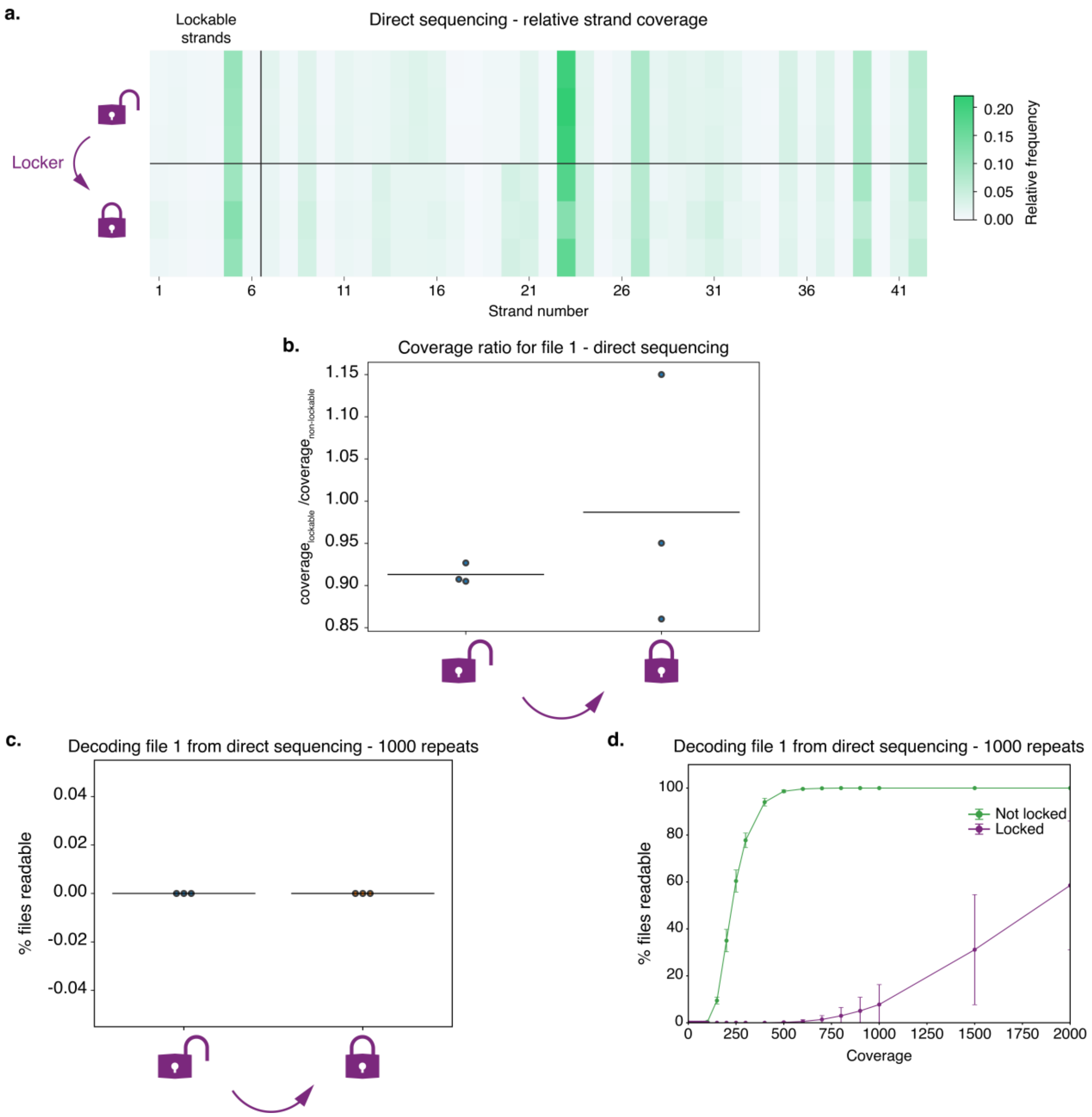

***Supplementary Figure 2 Attempted direct sequencing of locked File 1 a.*** *Heatmap of relative number of sequencing reads of file-encoding strands of File 1 from direct sequencing in not locked (top) and locked (bottom) state (Methods). Strands with a L region are indicated as "lockable strands". Each row shows relative sequence reads for one physical replicate, normalized to total reads from that replicate (n = 3).* ***b.*** *Ratio of relative File 1 reads for lockable strands to non-lockable strands in not locked and locked states. File samples were sequenced directly, and the relative number of reads was determined for each strand (Methods).* ***c.*** *Percentage of tries File 1 is readable from direct sequencing after 1000 simulated samplings at a fixed coverage of 30x. The percentage of trials File 1 could be decoded for each sequenced experiment was determined before locking (left) and in a locked state (right). In both states, File 1 could be decoded in 0% of the trials. Solid lines denote mean percentage of readable trials; points indicate individual experiments.* ***d.*** *Evaluating required sequencing coverage for reading File 1 from direct sequencing. 1000 simulated sampling were performed per sample per coverages ranging from 0 to 2000x. Points denote mean percentage of readable trials; whiskers denote standard deviations.*

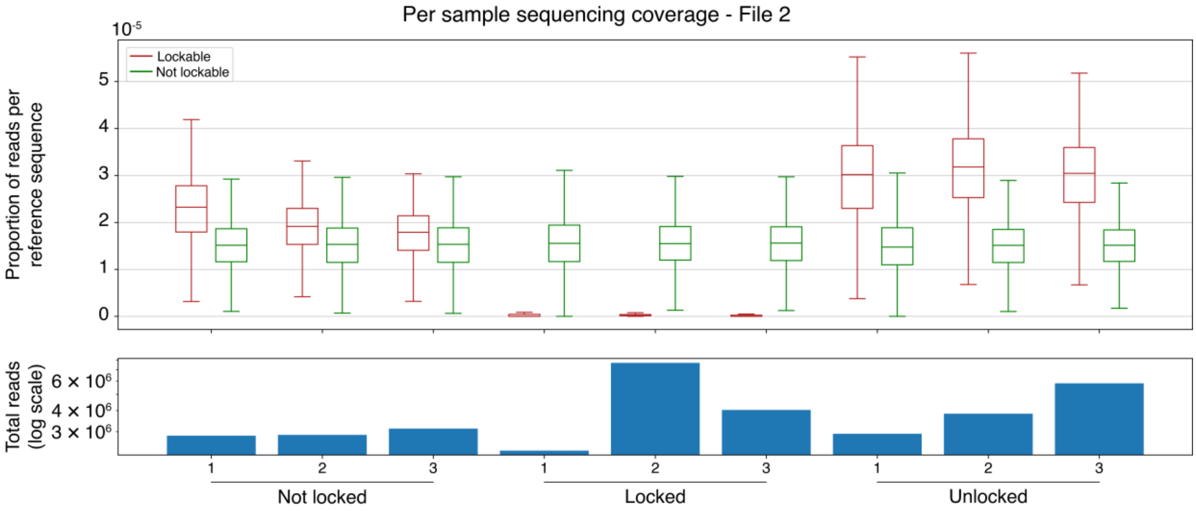

***Supplementary Figure 3 Coverage distributions for each experiment in Figure 4b*** *Sequencing coverage across independent experiments for each file state (Not locked, Locked and Unlocked). Top: Boxplots indicate coverage distribution for reads that were uniquely mapped to a single reference sequence. Lockable and non-lockable strands are shown as individual populations. Bottom: Total number of aligned reads per sample.*

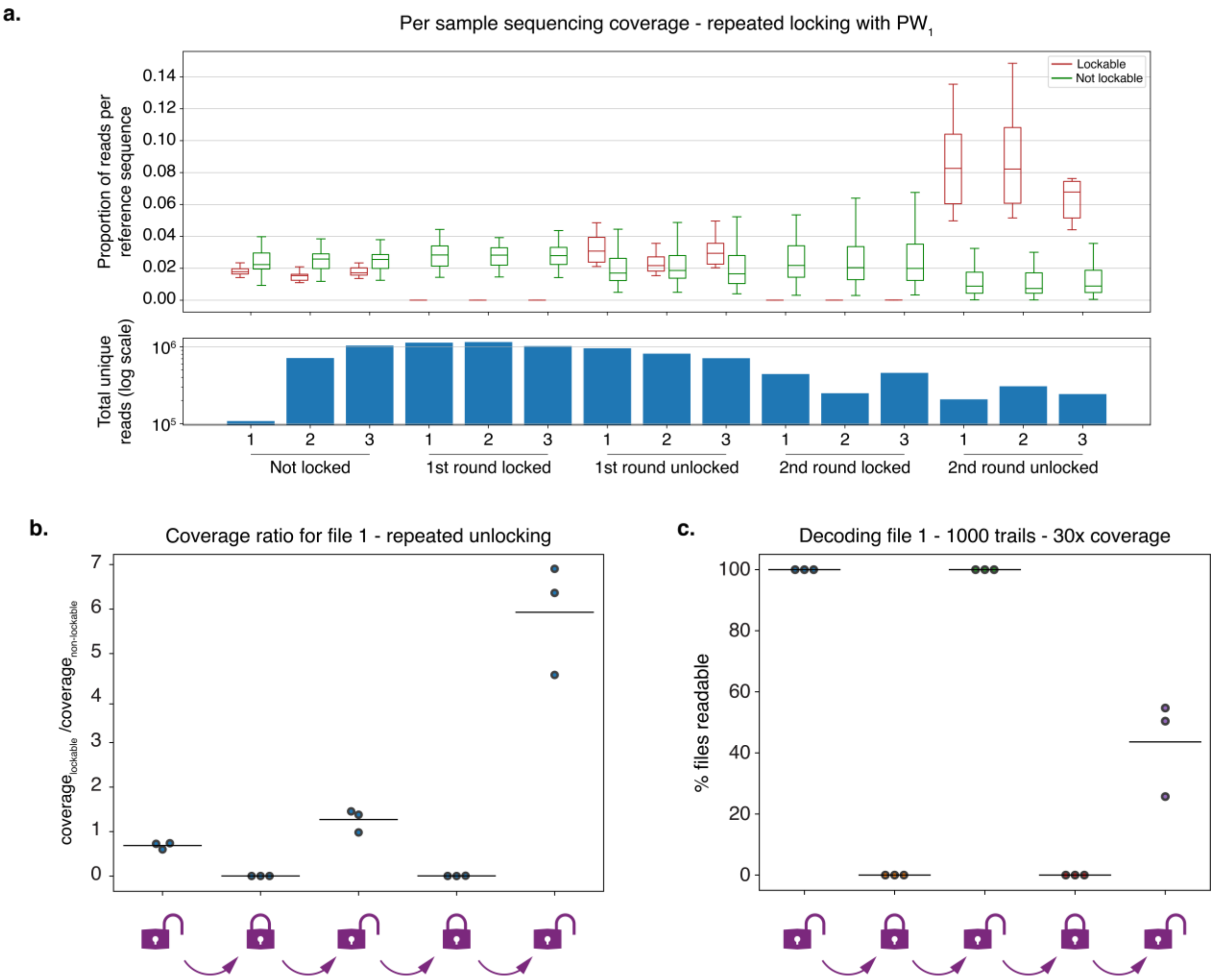

***Supplementary Figure 4 Decreased repeated locking performance unprotected key strand a.*** *Coverage distributions of separate DNA-GUARD locking and unlocking experiments for File 1 after two rounds of locking and unlocking using* ***$L_1$*** *and* ***$Key_1$****. Sequencing coverage across independent experiments for each file state (Not locked, 1st round locked, 1st round unlocked, 2nd round locked and*

*2nd round unlocked). Top: Boxplots indicate coverage distribution for reads that were uniquely mapped to a single reference sequence. Lockable (red) and non-lockable (green) strands are shown as individual populations. Bottom: Total number of aligned reads per sample.* ***b.*** *Ratio of relative File 1 reads for lockable strands to non-lockable strands in not locked, and both locked and unlocked states. PCR products were sequenced, and the relative number of reads was determined for each strand (Methods).* ***c.*** *Simulation used to determine functionality of DNA-GUARD after repeated locking and unlocking of File 1. Reads obtained from the locked PCR of each file are randomly sampled to simulate an average coverage of 30 reads per strand. Decoding is attempted, and the result is recorded. These steps are repeated 1000 times to determine the percentage of times File 1 can be accessed in each state. Solid lines denote mean percentage of readable trials; points indicate individual experiments. All experiments in the first locked state show 0% readability, and the not locked and 1st round unlocked state showed 100% readability. For the second round locked state, readability was 0.2% (0.0%, 0.1% and 0.5%) and in second round unlocked, mean readability dropped to 43.6% (50.4%, 25.7% and 54.7%).*

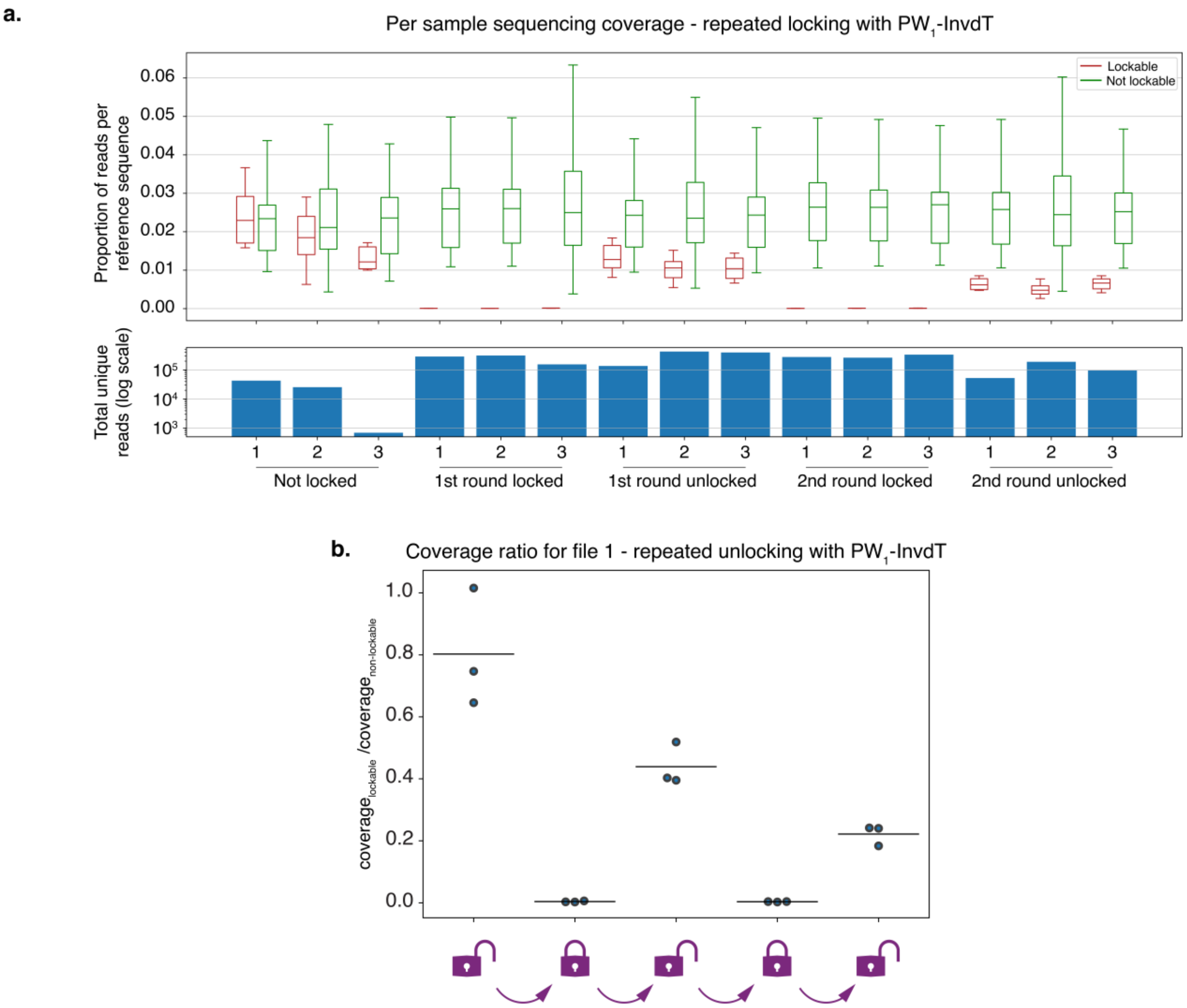

***Supplementary Figure 5 Coverage distributions for each experiment in Figure 4d a.*** *Coverage distributions of separate DNA-GUARD locking and unlocking experiments for File 1 after two rounds of locking and unlocking using* ***$L_1$*** *and* ***$Key_1$-InvdT****. Sequencing coverage across independent experiments for each file state (Not locked, 1st round locked, 1st round unlocked, 2nd round locked and 2nd round unlocked). Top: Boxplots indicate coverage distribution for reads that were uniquely mapped to a single reference sequence. Lockable (red) and non-lockable (green) strands are shown as individual populations. Bottom: Total number of aligned reads per sample.* ***b.*** *Ratio of relative File 1 reads for lockable stand to non-lockable strands in not locked, and both locked and unlocked states. PCR products were sequenced, and the relative number of reads was determined for each strand (Methods).*

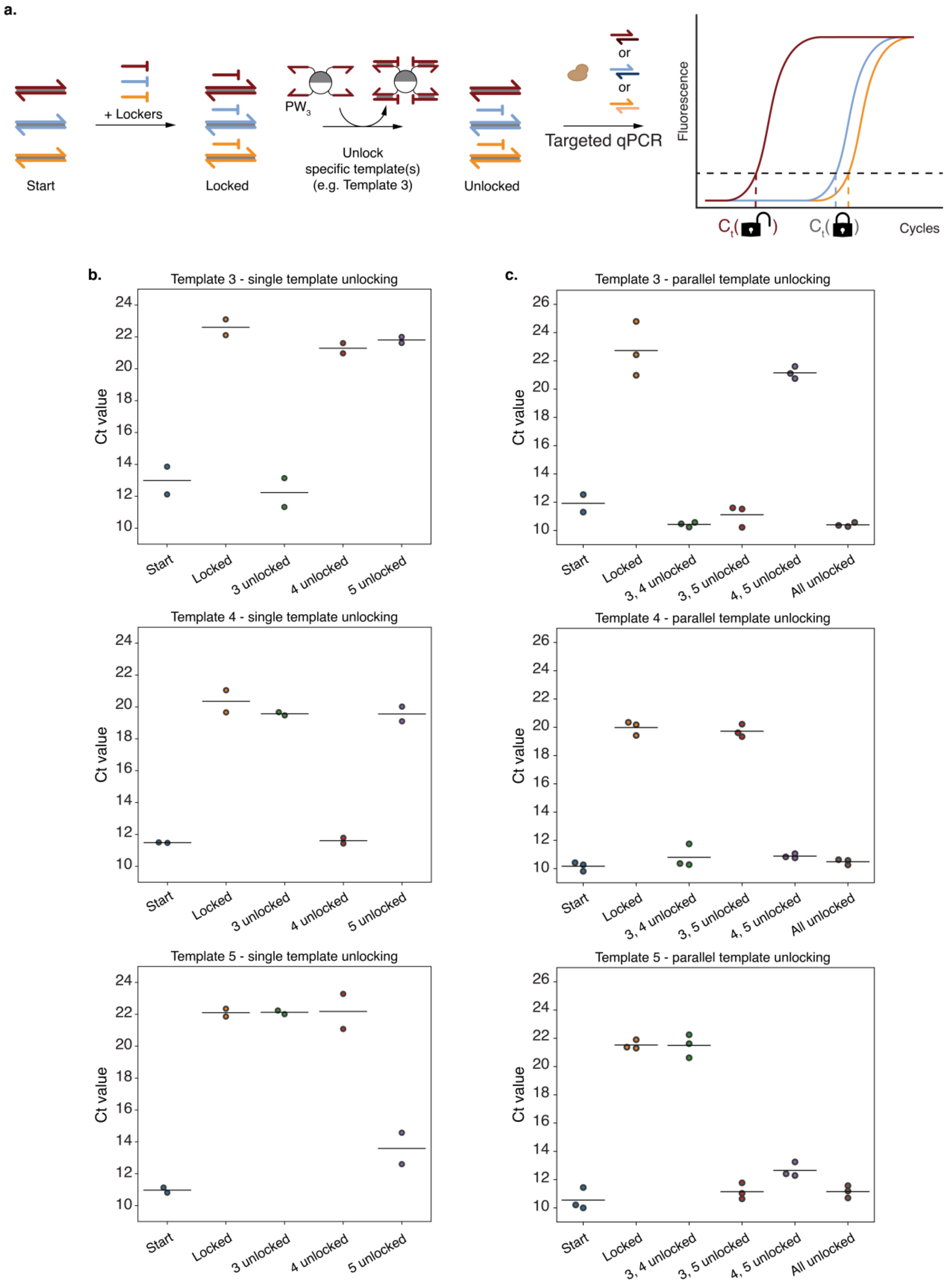

***Supplementary Figure 6 Functionality of orthogonally designed primer and locker combinations a.*** *Orthogonality test for **Key$_3$**, **Key$_4$** and **Key$_5$**. Lockable **Templates 3, 4 and 5** (red, blue and orange*

*respectively) were pooled together (Start) and all cognate lockers (**$L_3$**, **$L_4$** and **$L_5$**) were added (Locked). The combined locked pool was unlocked with either **$Key_3$**, **$Key_4$** and/or **$Key_5$** (**$Key_3$** is shown). Samples from each state were directly used in qPCR with primers for Template 3, 4 or 5. **b.** Ct values from qPCR results for template 3 (top), 4 (middle) and 5 (bottom) from single template unlocking experiments. N = 2, physical replicates. **c** Ct values from qPCR results for template 3 (top), 4 (middle) and 5 (bottom) from parallel template unlocking experiments. N = 3, physical replicates. Sequences for **Template 3**, **$FW_3$**, **$RV_3$**, **$L_3$**, and **$Key_3$**, for **Template 4**, **$FW_4$**, **$RV_4$**, **$L_4$**, and **$Key_4$** and for **Template 5**, **$FW_5$**, **$RV_5$**, **$L_5$**, and **$Key_5$** are given in Supplementary Table 9.*

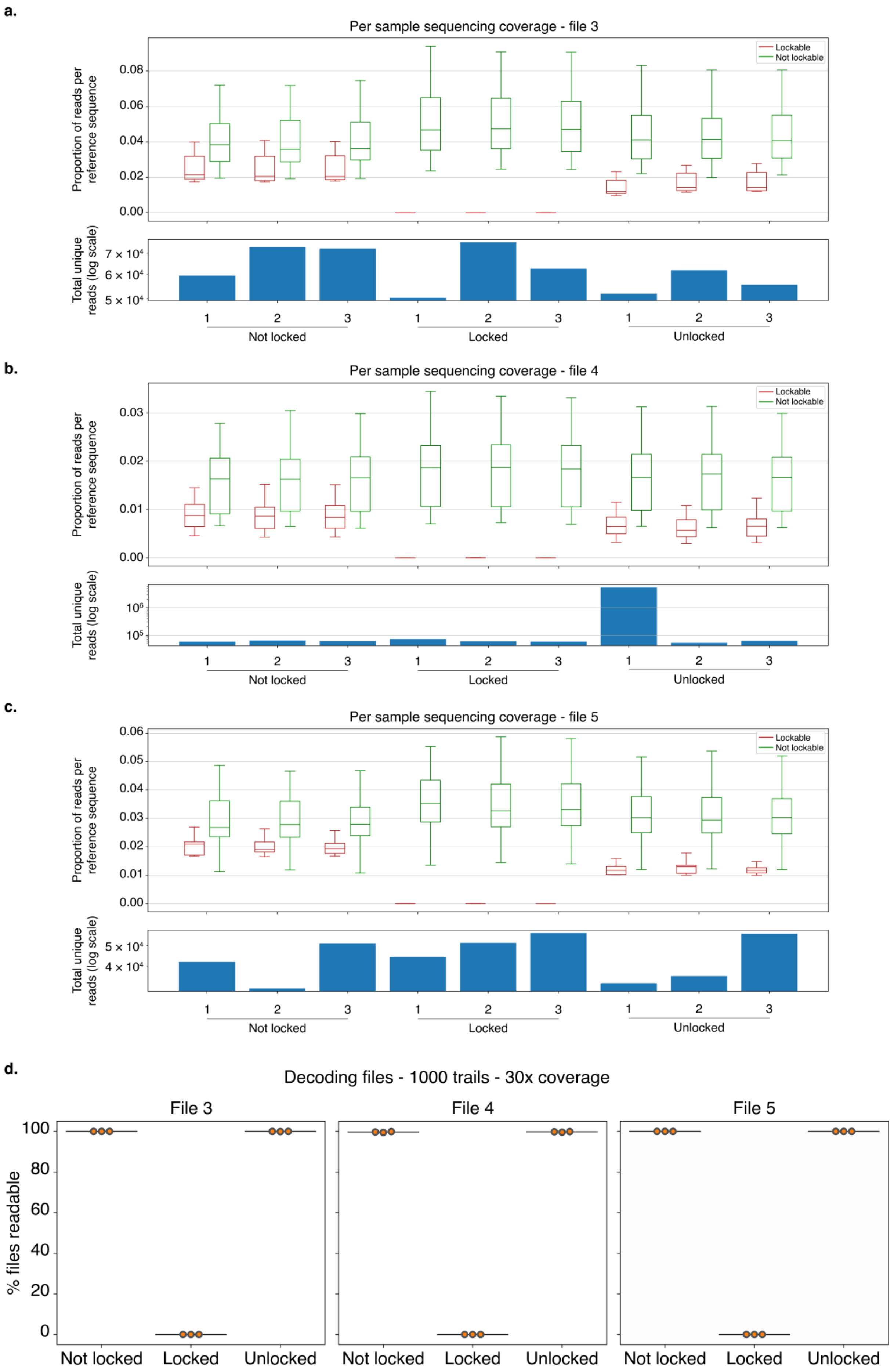
a.
Per sample sequencing coverage - file 3
Lockable
Not lockable
Proportion of reads per reference sequence
Total unique reads (log scale)
Not locked
Locked
Unlocked
b.
Per sample sequencing coverage - file 4
c.
Per sample sequencing coverage - file 5
d.
Decoding files - 1000 trails - 30x coverage
File 3
File 4
File 5
% files readable

***Supplementary Figure 7 DNA-GUARD efficiency for File 3, 4 and 5 a. – c.*** *Coverage distributions of separate DNA-GUARD locking and unlocking experiments for files 3 (**a.**), 4 (**b.**) and 5 (**c.**) used in Figure 5. Sequencing coverage across independent experiments for each file state (Not locked, Locked and Unlocked). Top: Boxplots indicate coverage distribution for reads that were uniquely mapped to a single reference sequence. Lockable (red) and non-lockable (green) strands are shown as individual populations. Bottom: Total number of aligned reads per sample.* ***d.*** *Simulation used to determine functionality of generated lockers for Files 3 – 5. Reads obtained from the locked PCR of each file are randomly sampled to simulate an average coverage of 30 reads per strand. Decoding is attempted, and the result is recorded. These steps are repeated 1000 times to determine the percentage of times each File can be accessed in each state. Results for File 3 (left), 4 (middle), and 5 (right) are plotted separately. Solid lines denote mean percentage of readable trials; points indicate individual experiments. All locked state samples for each of the files showed 0% readability. All not locked and unlocked state for file 3 and 5 showed 100% readability, with the exception of a single replicate for each in the unlocked state, both 99.9%. For file 4, there was 99.5%, 99.9% and 99.7% readability in the not locked state and 99.6%, 99.8% and 99.9% for the unlocked state.*

### Supplementary Tables

***Supplementary Table 1 – DNA sequences used in Figure 2, 3 and 4***

| **Name** | **Sequence** | **Length (# bases)** | **Modification** |
|---|---|---|---|
| $A_1$ | TGTCTCACTAGGTCCTCTCAGCTCTTCCCACCACCATTGCCTTC<br>TTCTTCCTTCTGAGTCTAACCACCACCATTGCCTTATCTACCAT<br>TCTGAGTCTTACCACCACCATTGCCATTTCATTATCTACGAGTC<br>TATCCACCACCATTGCAGGAGTTTCAGTTTCGACGC | 168 | |
| $A_2$ | GCGTCGAAACTGAAACTCCTGCAATGGTGGTGGATAGACTCGTA<br>GATAATGAAATGGCAATGGTGGTGGTAAGACTCAGAATGGTAGA<br>TAAGGCAATGGTGGTGGTTAGACTCAGAAGGAAGAAGAAGGCAA<br>TGGTGGTGGGAAGAGCTGAGAGGACCTAGTGAGACA | 168 | |
| $B_1$ | TGTCTCACTAGGTCCTCTCATTCCGAGGGAGATCTACACTGCTC<br>GTCTTAATATACGGAATAACCAGCCCGCGGCTCCAAACCGACTA<br>GATTCTTTCTCGAGTTATCCACATACATCGGTCTTATATTGTTG<br>GTGATACCCAAACAATAGGAGTTTCAGTTTCGACGC | 168 | |
| $B_2$ | GCGTCGAAACTGAAACTCCTATTGTTTGGGTATCACCAACAATA<br>TAAGACCGATGTATGTGGATAACTCGAGAAAGAATCTAGTCGGT<br>TTGGAGCCGCGGGCTGGTTATTCCGTATATTAAGACGAGCAGTG<br>TAGATCTCCCTCGGAATGAGAGGACCTAGTGAGACA | 168 | |
| $U_{FW}$ | TGTCTCACTAGGTCCTCTCA | 20 | |
| $U_{RV}$ | GCGTCGAAACTGAAACTCCT | 20 | |
| $F_A$ | CCACCATTGCCTTCTTCTTC | 20 | |
| $R_A$ | GGATAGACTCGTAGATAATG | 20 | |
| $F_B$ | CCGAGGGAGATCTACACTGC | 20 | |
| $R_B$ | GTATCACCAACAATATAAGAC | 21 | |
| $L_1$ | CACTAGGTCCTCTCAGCTCTTCCCACCACCATTGCCTTCTTCT | 43 | 3’ inverted dT |
| $Key_1$ | AAGAAGAAGGCAATGGTGGTGGGAAGAGCTGAGAGGACCTAGTG | 44 | 5’ biotin |
| $Key_S$ | TCGTCGTTGATTGGTATAGGCCCTTCAATCCGCGAACCAGAGCT | 44 | 5’ biotin |
| $Key_1$-InvdT | AAGAAGAAGGCAATGGTGGTGGGAAGAGCTGAGAGGACCTAGTG | 44 | 5’ biotin<br>3’ inverted dT |

***Supplementary Table 2 – Ct values figure 2c***

| | **A** | **B** | **ΔCt(A,B)** |
|---|---|---|---|
| **0 nM** | 10.25 | 9.36 | 0.89 |
| | 9.31 | 8.62 | 0.69 |
| | 9.27 | 8.25 | 1.02 |
| **50 nM** | 10.37 | 8.73 | 1.64 |
| | 10.66 | 8.44 | 2.22 |
| | 10.71 | 8.23 | 2.48 |
| **500 nM** | 17.28 | 8.62 | 8.66 |
| | 17.98 | 8.44 | 9.54 |
| | 17.53 | 8.34 | 9.19 |
| **5000 nM** | 18.74 | 8.91 | 9.83 |
| | 18.61 | 8.51 | 10.1 |
| | 18.21 | 8.26 | 9.95 |

*Supplementary Table 3 – Ct values figure 2e*

| | A | B | ΔCt(A,B) |
|---|---|---|---|
| **Not locked** | 15.14 | 14.18 | 0.96 |
| | 13.94 | 12.17 | 1.76 |
| | 14.57 | 12.23 | 2.34 |
| **Locked** | 23.93 | 12.24 | 11.69 |
| | 24.26 | 12.32 | 11.94 |
| | 24.76 | 12.56 | 12.20 |
| **Unlocked** | 15.29 | 12.12 | 3.17 |
| | 16.48 | 12.37 | 4.11 |
| | 17.31 | 16.21 | 1.10 |

*Supplementary Table 4 – Files encoded in DNA*

| File # | File name | Origin | Contents |
|---|---|---|---|
| 1 | The_crazy_ones.txt | Apple "Think Different" advertisement | Text: "Here's to the crazy ones. The misfits. The rebels. The troublemakers. The round pegs in the square holes. The ones who see things differently. They're not fond of rules. And they have no respect for the status quo. You can quote them, disagree with them, glorify or vilify them. About the only thing you can't do is ignore them. Because they change things. They push the human race forward. And while some may see them as the crazy ones, we see genius. Because the people who are crazy enough to think they can change the world, are the ones who do." |
| 2 | Operation_overlord.pdf | Supreme headquarters, Allied expeditionary force (1944) | 14 pages of the official command directive for the invasion of northwest Europe by the allied forces during the second world war. |
| 3 | Frankenstein.txt | "Frankenstein or The Modern Prometheus" by Mary Shelley (1818) | Text: "None but those who have experienced them can conceive of the enticements of science. In other studies you go as far as others have gone before you, and there is nothing more to know; but in scientific pursuit there is continual food for discovery and wonder." |
| 4 | Molecular_structure_of_nucleic_acids.txt | "Molecular Structure of Nucleic Acids: A | Text: "If it is assumed that the bases only occur in the structure in the most plausible tautomeric forms (that is, with |

| | | | |
|---|---|---|---|
| | | Structure for Deoxyribose Nucleic Acid" by J.D. Watson and F.H.C. Crick (1953) | the keto rather than the enol configurations) it is found that only specific pairs of bases can bond together. These pairs are : adenine (purine) with thymine (pyrimidine), and guanine (purine) with cytosine (pyrimidine). In other words, if an adenine forms one member of a pair, on either chain, then on these assumptions the other member must be thymine ; similarly for guanine and cytosine. The sequence of bases on a single chain does not appear to be restricted in any way. However, if only specific pairs of bases can be formed, it follows that if the sequence of bases on one chain is given, then the sequence on the other chain is automatically determined. " |
| 5 | Origin_of_species.txt | *"On the Origin of Species by Means of Natural Selection, or the Preservation of Favoured Races in the Struggle for Life"* by Charles Darwin (1859) | Text: "The resemblance of the offspring to its parents—the retention of peculiarities, and the reappearance of long-lost characters—can, I think, be truly said to be due to the transmission of a something which, though at present unknown, must be material in its nature. It must be a substance which can be added to or modified, but which, when once formed, remains persistent for many generations." |

***Supplementary Table 5 – Strand composition parameters DNA-encoded files***

| **File** | **File strands** | **Lockable strands** | **Tolerated loss of lockable strands** | **Required lockable strands** | **Lockable fraction** |
|---|---|---|---|---|---|
| 1 | 42 | 6 | 4 | 2 | 14.3% |
| 2 | 65,419 | 1,200 | 1,100 | 100 | 1.8% |
| 3 | 28 | 8 | 6 | 2 | 28.6% |
| 4 | 68 | 12 | 10 | 2 | 17.6% |
| 5 | 38 | 9 | 7 | 2 | 23.7% |

***Supplementary Table 6 – File 1 coverage ratio between lockable strands and non-lockable strands used in Figure 3d***

| | **Mean reads per lockable strand** | **Mean reads per non-lockable strand** | **Ratio lockable to non-lockable strands** |
|---|---|---|---|
| **Not locked** | 20887 | 30373 | 0.688 |
| | 20540 | 30547 | 0.672 |
| | 18675 | 27484 | 0.679 |

| | | | |
|---|---|---|---|
| **Locked** | 90.3 | 30858 | 0.00292 |
| | 80.5 | 34185 | 0.00235 |
| | 62.3 | 19929 | 0.00313 |
| **Unlocked** | 25836 | 19760 | 1.307 |
| | 24967 | 24953 | 1.001 |
| | 30393 | 23708 | 1.282 |
| **Scrambled Key** | 135.8 | 32134 | 0.00423 |
| | 129.5 | 34802 | 0.00372 |
| | 314.7 | 30721 | 0.0102 |

***Supplementary Table 7 – File 1 readability results for repeated unlocking with Key$_1$ (1000 repeats per replicate) used in Supplementary Figure 4c***

| | **Sample #1** | **Sample #2** | **Sample #3** |
|---|---|---|---|
| **Not locked** | 100 | 100 | 100 |
| **1st round locked** | 0.0 | 0.0 | 0.0 |
| **1st round unlocked** | 100 | 100 | 100 |
| **2nd round locked** | 0.0 | 0.1 | 0.5 |
| **2nd round unlocked** | 50.4 | 25.7 | 54.7 |

***Supplementary Table 8 – File 1 readability results for repeated unlocking with Key$_1$-InvdT (1000 repeats per replicate) used in Figure 4d***

| | **Sample #1** | **Sample #2** | **Sample #3** |
|---|---|---|---|
| **Not locked** | 99.1 | 97.3 | 91.3 |
| **1st round locked** | 0.0 | 0.1 | 0.1 |
| **1st round unlocked** | 99.1 | 96 | 97.7 |
| **2nd round locked** | 0.1 | 0.1 | 0.0 |
| **2nd round unlocked** | 97.2 | 82.5 | 94.4 |

***Supplementary Table 9 – DNA sequences used in Figure 5 and Supplementary Figure 5 and 6***

| **Name** | **Sequence** | **Length (# bases)** | **Modification** |
|---|---|---|---|
| Template 3 | AACATCGTGTCCAAGCAAGTTCTCCAGCAGTCCTTCAATA CAGCCATTCTGCTTGCGTCGCTGCTGCTGCTGTCACGGAT GCACGTCTACAGGCTGCTGCTCGTACATGTTGATTGTTTG TCCACGCTTTCGA | 133 | |
| FW$_3$ | AACATCGTGTCCAAGCAAGT | 20 | |
| RV$_3$ | TCGAAAGCGTGGACAAACAA | 20 | |
| L$_3$ | CGTGTCCAAGCAAGTTCTCCAGCAGTCCTTCAATACAGCC ATTC | 44 | 3' inverted dT |
| Key$_3$ | AGAATGGCTGTATTGAAGGACTGCTGGAGAACTTGCTTGG ACACG | 45 | 5' biotin |
| Template 4 | TTAATCGGTAACACCTGCGGATTTCACTCTGTCTCATGCC TCTCTACGTTGCTCGAGTCACTGCTGCTGCTAGAACACAG GCAGTCTGCCTGGCTGCTGCTTGTATGTAGGAGATTGTTA CGAATCGGTGCCA | 133 | |
| FW$_4$ | TTAATCGGTAACACCTGCGG | 20 | |
| RV$_4$ | TGGCACCGATTCGTAACAAT | 20 | |

| | | | |
|---|---|---|---|
| $L_4$ | CGGTAACACCTGCGGATTTCACTCTGTCTCATGCCTCTCTACGT | 44 | 3’ inverted dT |
| $Key_4$ | AACGTAGAGAGGCATGAGACAGAGTGAAATCCGCAGGTGTTACCG | 45 | 5’ biotin |
| Template 5 | TGTGTTCCTCCTCGGTATGATAGCCTGTGACTCGACTTCTGAACCTACTTGCTCGATACGCTGCTGCTGCTCTAGTCCGAACAACTTGCGCTGCTGCTACTATGCATTCAAGGAAGCGCCAACTAATTGT | 133 | |
| $FW_5$ | TGTGTTCCTCCTCGGTATGA | 20 | |
| $RV_5$ | ACAATTAGTTGGCGCTTCCT | 20 | |
| $L_5$ | TCCTCCTCGGTATGATAGCCTGTGACTCGACTTCTGAACCTACT | 44 | 3’ inverted dT |
| $Key_5$ | AAGTAGGTTCAGAAGTCGAGTCACAGGCTATCATACCGAGGAGGA | 45 | 5’ biotin |

***Supplementary Table 10 – File 3 readability results for parallel unlocking with $Key_3$, $Key_4$ and $Key_5$. (1000 repeats per replicate) used in Figure 5e***

| State | Sample #1 | Sample #2 | Sample #3 |
|---|---|---|---|
| **Not locked** | 100.0 | 100.0 | 100.0 |
| **All locked** | 0.0 | 0.0 | 49.9 |
| **File 3 unlocked** | 100.0 | 100.0 | 99.7 |
| **File 4 unlocked** | 0.0 | 0.0 | 0.0 |
| **File 5 unlocked** | 0.0 | 0.0 | 0.0 |
| **File 3 and 4 unlocked** | 100.0 | 100.0 | 99.9 |
| **File 3 and 5 unlocked** | 100.0 | 100.0 | 99.7 |
| **File 4 and 5 unlocked** | 0.0 | 0.0 | 1.3 |
| **File 3, 4 and 5 unlocked** | 100.0 | 100.0 | 100.0 |

***Supplementary Table 11 – File 4 readability results for parallel unlocking with $Key_3$, $Key_4$ and $Key_5$. (1000 repeats per replicate) used in Figure 5e***

| State | Sample #1 | Sample #2 | Sample #3 |
|---|---|---|---|
| **Not locked** | 100.0 | 100.0 | 99.8 |
| **All locked** | 0.0 | 0.0 | 0.0 |
| **File 3 unlocked** | 31.3 | 0.0 | 0.0 |
| **File 4 unlocked** | 100.0 | 100.0 | 77.0 |
| **File 5 unlocked** | 0.0 | 0.0 | 0.0 |
| **File 3 and 4 unlocked** | 100.0 | 100.0 | 51.0 |
| **File 3 and 5 unlocked** | 0.0 | 0.0 | 0.0 |
| **File 4 and 5 unlocked** | 100.0 | 99.9 | 100.0 |
| **File 3, 4 and 5 unlocked** | 100.0 | 100.0 | 100.0 |

***Supplementary Table 12 – File 5 readability results for parallel unlocking with $Key_3$, $Key_4$ and $Key_5$. (1000 repeats per replicate) used in Figure 5e***

| State | Sample #1 | Sample #2 | Sample #3 |
|---|---|---|---|
| **Not locked** | 100.0 | 100.0 | 100.0 |
| **All locked** | 0.0 | 0.0 | 1.2 |
| **File 3 unlocked** | 0.0 | 0.0 | 0.0 |
| **File 4 unlocked** | 0.0 | 0.0 | 0.0 |

| | | | |
|---|---|---|---|
| **File 5 unlocked** | 100.0 | 100.0 | 99.5 |
| **File 3 and 4 unlocked** | 0.0 | 0.0 | 0.2 |
| **File 3 and 5 unlocked** | 100.0 | 100.0 | 99.8 |
| **File 4 and 5 unlocked** | 100.0 | 100.0 | 99.7 |
| **File 3, 4 and 5 unlocked** | 100.0 | 100.0 | 99.5 |
